# TORC1 spatial segregation and hysteresis are regulated by the small GTPase Arf1

**DOI:** 10.64898/2026.04.29.721593

**Authors:** Ludovic Enkler, Trian Nuredini, Aleksei Mironov, Danilo Ritz, Congwei Wang, Mihaela Zavolan, Anne Spang

## Abstract

TORC1 is a central regulator of growth, integrating inputs from kinases and signaling proteins. How TORC1 activity is regulated, is still not completely understood. The small GTPase Arf1 has been implicated in TORC1 regulation, but the underlying mechanism remains elusive. Here, we show that Arf1 controls TORC1 in an allele-specific manner under nutrient and heat stress in yeast. Unexpectedly, both the hyperactive mutant *arf1-11* and the loss-of-function mutant *arf1-18* reduce TORC1 activity, while other loss-of-function alleles did not. We reveal two distinct functions of Arf1: first, it ensures Golgi-to-vacuole/lysosome transport for the functional maintenance of these organelles; second, it regulates TORC1 activity at the ER-Golgi-vacuole interface. Hyperactive Arf1-11 drives sequestration of Kog1 and Tor1 into cytoplasmic foci, to which Arf1-11 can be equally recruited. Importantly, Arf1 also promotes TORC1 re-activation during stress recovery - so-called hysteresis. Our data reveal an unexpected dual function for Arf1 as a negative and a positive regulator of TORC1 activity in a context-dependent manner.

**Teaser:** The small GPTase Arf1 acts as a dual switch for the key cellular growth regulator TORC1 by increasing or decreasing TORC1 activity depending on nutritional status and stress.

## Introduction

Continuous adaptation to fluctuating environmental and intracellular conditions is key to maintaining metabolic homeostasis. Central to this adaptive capacity is the Target of Rapamycin Complex 1 (TORC1), a highly conserved protein kinase complex, which integrates signals related to nutrient availability, energy status, and cellular stress (reviewed in 1–3). By modulating a wide range of anabolic and catabolic processes, TORC1 serves as a master regulator of cell growth and division. While the spatial and temporal dynamics of TORC1 have emerged as a key aspect of its function, many fundamental aspects of TORC1 regulation remain elusive.

Small GTPases of the ADP-ribosylation factor (Arf) family play a crucial role in regulating intracellular traffic, organelle dynamics, and cellular signaling. Among them, Arf1 is a key mediator of vesicle formation and membrane remodeling, and its activity is modulated by ArfGAPs (GTPase activating proteins) and ArfGEFs (guanine nucleotide exchange factors) which stimulate the GTP hydrolysis and the GDP-to-GTP exchange, respectively. Emerging evidence indicates that Arf1 functions beyond vesicle trafficking by regulating mRNA transport^4,5^, mitochondrial division and homeostasis^6–8^ as well as fatty acid storage and utilization^9–11^. Additionally, Arf1 regulates TORC1 in response to glutamine levels independently of the Rag guanosine triphosphatases (GTPases) normally required to recruit and activate mTORC1 ^12,13^, and its GTP-bound form counteracts S6K1 phosphorylation by mTOR^14^. Recent studies have also linked COPI vesicle biogenesis to mTORC1 activation and its relocation from the ER to the lysosome upon amino acid stimulation^15^. In mammals, the GTPase-activating protein ArfGAP1 has been found to sequester mTORC1 away from the lysosome, preventing its activation by the ras-like GTPase Rheb in the absence of amino acids^16^, reinforcing the idea that mTORC1 activity is closely linked to its localization on the lysosome/vacuole^17^.

Recent studies have highlighted the formation of specialized structures, such as TOROIDS and TOR bodies, which are induced under nutrient stress or energy depletion^18–20^. These structures are thought to facilitate the sequestration and inactivation of TORC1 and represent a pool of TORC1 available for fast reactivation after stress release, so-called hysteresis. Despite the recognized importance of these TORC1 sequestering structures, the molecular mechanisms and upstream regulators driving their formation remain poorly understood. Moreover, whether Arf1 contributes to the dynamic reorganization of TORC1 into TOROIDS or TOR bodies, and how this process intersects with the broader metabolic and stress signaling landscape, remain unexplored.

In this study, we identify a previously unrecognized link between Arf1, TORC1 activity and the promotion TORC1 sequestering structures. We show that Arf1 activity modulates TORC1 activity in a GAP- and GEF-dependent manner. Using temperature-sensitive (*ts*) mutants of Arf1 and mutants in ArfGAPs and ArfGEFs, we demonstrate that Arf1 activity influences TORC1 sequestering structures. Those structures harbor both the Kog1 regulatory subunit and the Tor1 kinase and are found in the cytoplasm close to the ER of yeast cells in response to various stresses. More importantly, starvation-refeeding assays suggest that Arf1 activity is not only linked to sequester inactive TORC1 but also to TORC1 re-activation. Our findings reveal a critical intersection between Arf1 activity, nutrient and stress sensing, and TORC1 hysteresis. These insights provide a new perspective on the complex interplay between intracellular trafficking, membrane contact sites, nutrient signaling and stress, and shed light on a fundamental mechanism with potential implications for both physiology and disease.

## Results

### TORC1 activity is modulated by Arf1

To get a holistic and comprehensive view on how Arf1 impacts TORC1, we took an omics approach combining RNA sequencing, ribosome footprint sequencing and proteomics (**Fig. 1A**). We took advantage of the Arf1 hyperactive mutant strain *arf1-11* to investigate changes induced by Arf1 activity^10,21^. Among the most striking effects of Arf1 hyperactivity compared to wild-type ARF1 were the decrease of proteins involved in ribosomal RNA (rRNA) processing, ribosomal subunit processing, peptides biosynthetic process and actin filament reorganization. Those GO terms all point to global attenuation of translation (**Fig. 1B, 1C; Fig. S1A-D; Table S1-S3**). Down regulation of ribosome biosynthesis and translational attenuation are hallmarks of reduced TORC1 activity^22,23^ as well as actin regulation^24,25^.

**Figure 1.**
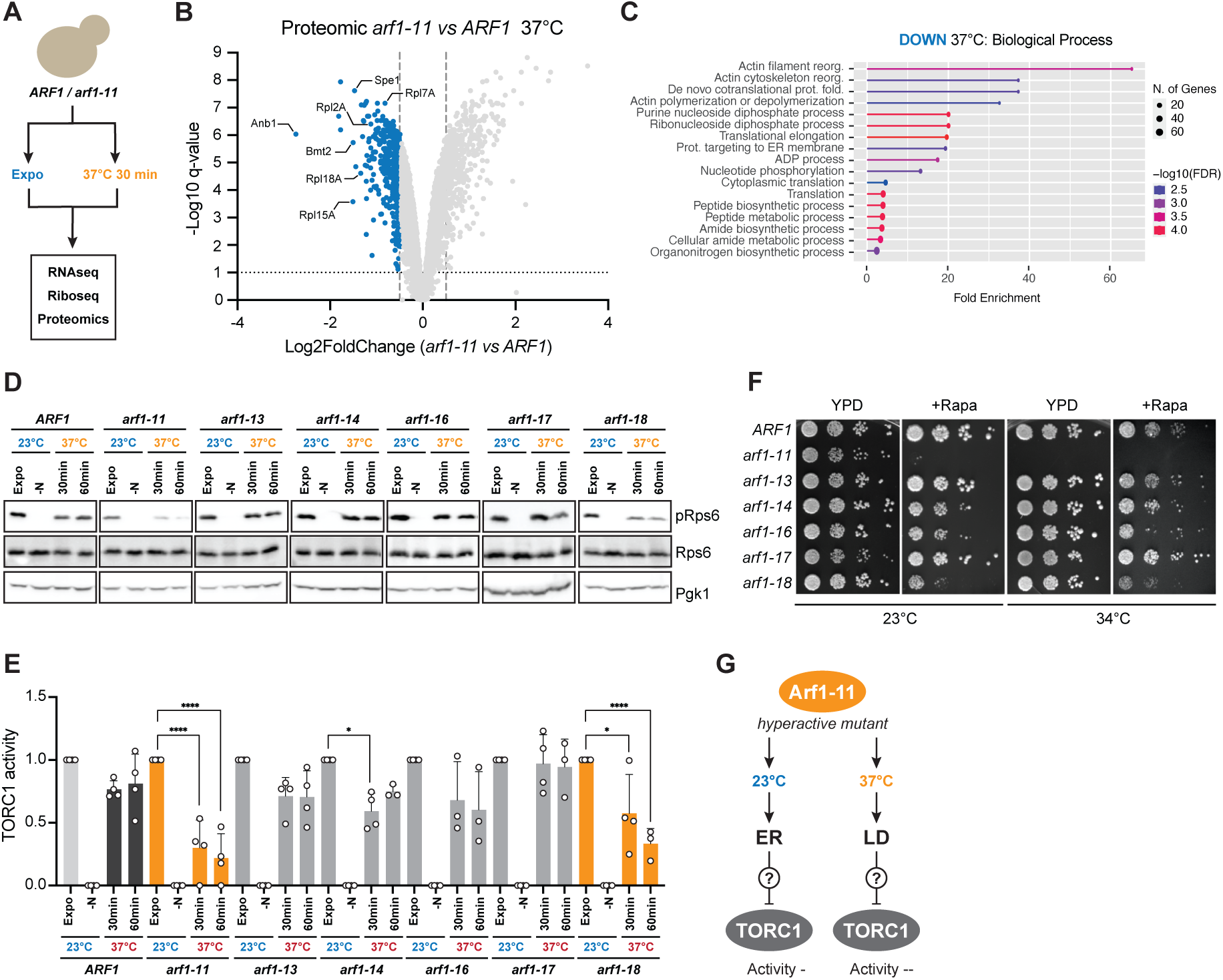
TORC1 activity is modulated by Arf1. **A)** Schematic depiction of the strategy used to identify pathways deregulated in the *arf1-11 ts*-allele mutant strain. **B)** Volcano plot obtained from *arf1-11 vs ARF1* strains proteomics. All decreased proteins in *arf1-11* at 37°C are shown in blue. A subset of decreased ribosomal proteins are shown. **C)** Gene Ontology terms associated to yeast biological processes based on decreased proteins shown in (**B**). See also Supplementary Table 1. **D)** Immunoblot of the ribosomal protein Rps6 and its phosphorylated form pRps6 were performed in 6 *ts*-allele mutants of *ARF1*, as well as WT *ARF1* strain. The phosphoglycerate kinase 1 (Pgk1) was used as loading control. Cells were grown to exponential phase at 23°C for 6 h (Expo), shifted for 1 h in media lacking nitrogen sources (-N), or shifted for 30 or 60 minutes to 37°C. **E)** Relative fold changes in protein levels from immunoblots in (**D)**. Mean and standard deviation are shown from at least n = 3 biological replicates. Two-way ANOVA with Turkey test was performed. *arf1-11* Expo vs *arf1-11* 30min \*\*\*\**P*: 0.000000544900853; *arf1-11* Expo vs *arf1-11* 60min \*\*\*\**P*: 0.000000016420113; *arf1-14* Expo vs *arf1-14* 30min \**P*: 0.033269079746617; *arf1-18* Expo vs *arf1-18* 30min \**P*: 0.019322355811631; *arf1-18* Expo vs *arf1-18* 60min \*\*\*\**P*: 0.000016687988456. **F)** Growth test of ARF1, *arf1-11, arf1-13, arf1-14, arf1-16, arf1-17* and *arf1-18* strains on rich medium (YPD), YPD+Rapamycin (5ng/mL) incubated at 23°C and 34°C. **G)** Schematic of the localization of Arf1-11 mutant at 23°C and 37°C in yeast, correlated with TORC1 activity measured in (**D**) and (**E**).

To better understand the role of Arf1 in TORC1 signaling, we used a series of 7 *ts*-mutants^21^ with allele-specific phenotypes, which allow us to distinguish between various intracellular trafficking defects^10^. We then correlated allele-specific defects to changes in TORC1 activity by following the phosphorylation of the ribosomal protein Rps6 on Ser232-233^21,26^. The different *ts*-mutants were shifted from exponential phase at 23°C (referred to as permissive temperature) to 37°C (non-permissive temperature) for 30 or 60 min. In WT cells, the temperature shift to 37°C for 30 min induced a slight decrease in TORC1 activity, which was restored within 60 min (**Fig. 1D**). TORC1 activity was already reduced in exponentially growing *arf1-11* cells at 23°C, and this effect was exacerbated at 37°C, confirming our omics data (**Fig. 1D, 1E**). Among the other mutants, TORC1 activity was only decreased in *arf1-18* cells at the non-permissive temperature (**Fig. 1D, 1E**). While we previously established that *arf1-11* is a hyperactive mutant^10^, *arf1-18* is considered to be a loss-of-function mutant^21^. Both mutant proteins were well expressed (**Fig. S1E, S1F**), suggesting that Arf1 cycling between its active and inactive state is necessary to modulate TORC1 activity. Our mutant analysis also highlights that not all Arf functions affect TORC1 activity. Arf1’s best established functions are at the Golgi in the generation of transport vesicles. However, we did not observe Golgi defects by light microscopy (**Fig. S1G**). Consistent with the activity assay, *arf1-11* and *arf1-18* strains were rapamycin-sensitive both at permissive and non-permissive temperature (**Fig. 1F)**. This led us to conclude that distinct *arf1* mutants lead to reduced TORC1 activity, and that cycling between the GTPase active and inactive state might be required to regulate TORC1 activity (**Fig. 1G**).

### Arf1 activity regulates TORC1 activity

Arf1 activity is strictly controlled by ArfGAPs and ArfGEFs. If our assumption was correct, we should be able to identify the ArfGEF and ArfGAP involved in TORC1 regulation. In yeast, the two ArfGAP Glo3 and Gcs1 act on the Golgi-to-ER retrograde pathway^27^, while the ArfGAP Age1 and Age2 perform their function on the *trans*-Golgi (TGN) side (**Fig. 2A**). The ArfGEF Gea1 mainly acts on the Golgi-to-ER side^28,29^, while Gea2 is mainly operating within the Golgi, i.e. at the *trans*-, *mid*- and *cis*-Golgi^28,30^ (**Fig. 2A**). The ArfGEF Sec7 functions on the *trans*-Golgi side to regulate trafficking towards the plasma membrane, endosomes and the vacuole^31^ (**Fig. 2A**). Overlapping functions for Glo3 and Gcs1 as well as Gea1 and Gea2 have been demonstrated^27,32^.

**Figure 2.**
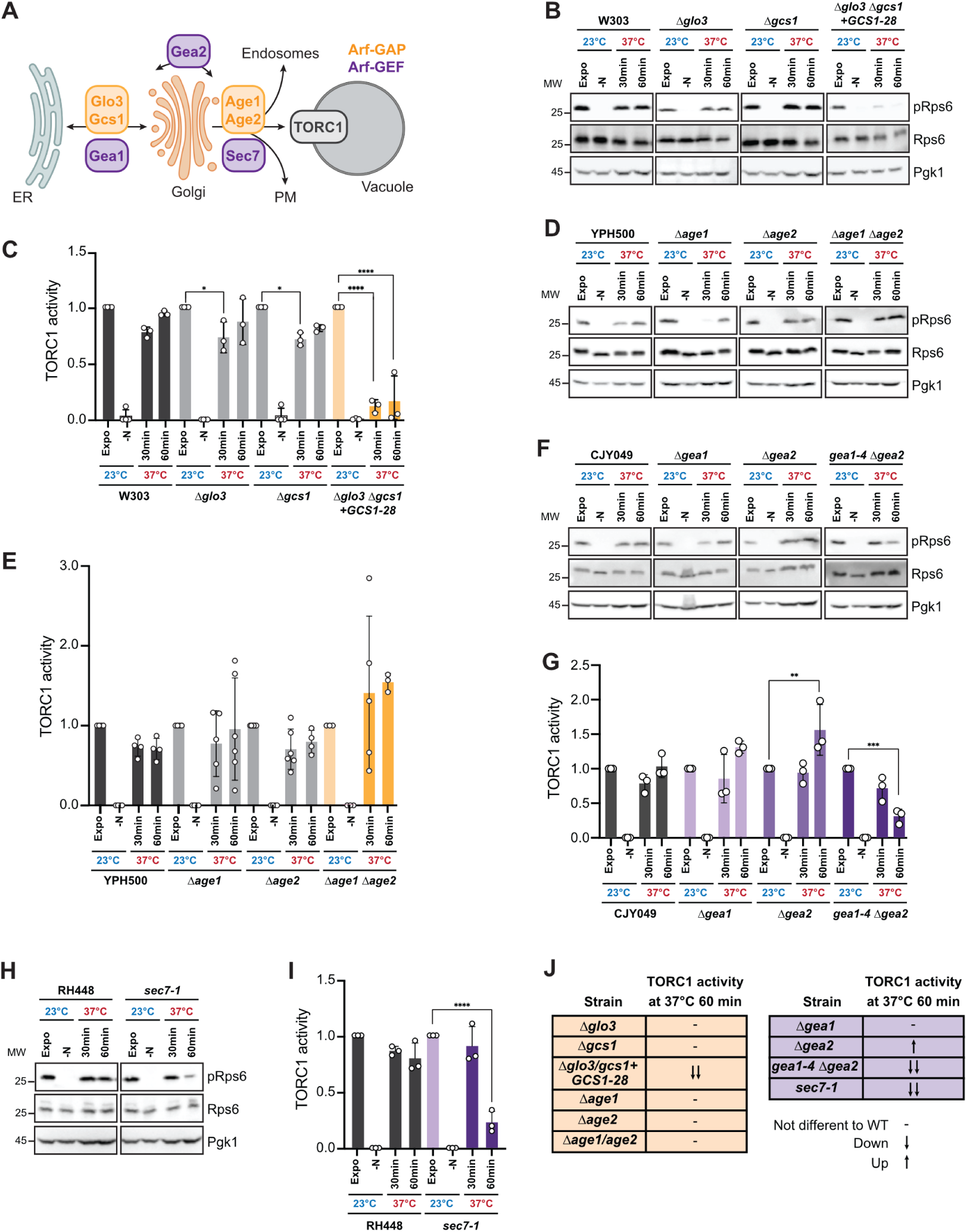
Arf1 activity regulates TORC1 activity. **A)** Schematic of ArfGAP (orange) and ArfGEF (purple) localization in cells with respect to ER, Golgi and the vacuole. PM: plasma membrane. **B)** Immunoblot of the ribosomal protein Rps6 and its phosphorylated form pRps6 were performed in W303 background strain, strains deleted for the ArfGAP *GLO3* (Δ*glo3*) or *GCS1* (Δ*gcs1*) or both rescued by a *ts*-allele of *GCS1* (Δ*glo3 Δgcs1+GCS1-28*). Pgk1 was used as loading control. Cells were grown to exponential phase at 23°C for 6 h (Expo), shifted for 1 h in media deprived of nitrogen sources (-N), or shifted for 30 or 60 minutes to 37°C. **C)** Relative TORC1 activity based on protein level changes from immunoblots performed in (**B**). Mean and standard deviation are shown from n = 3 biological replicates. Two-way ANOVA with Turkey test was performed. *Δglo3* Expo vs *Δglo3* 30min \**P*: 0.048012775181558; *Δgcs1* Expo vs *Δgcs1* 30min \**P*: 0.023761773769372; Δ*glo3 Δgcs1+GCS1-28* Expo *vs* Δ*glo3 Δgcs1+GCS1-28* 30min *\*\*\*\*P*<0.000000000000001; Δ*glo3 Δgcs1+GCS1-28* Expo *vs* Δ*glo3 Δgcs1+GCS1-28* 60min *\*\*\*\*P*<0.000000000000001. **D)** Immunoblot of the ribosomal protein Rps6 and its phosphorylated form pRps6 were performed in YPH500 background strain, strains deleted for the ArfGAP *AGE1* (Δ*age1*) or *AGE2* (Δ*age2*), or both (Δ*age1 Δage2*). Pgk1 was used as loading control. Cells were grown to exponential phase at 23°C for 6 h (Expo), shifted for 1 h in media deprived of nitrogen sources (-N), or shifted for 30 or 60 minutes to 37°C. **E)** Relative TORC1 activity based on protein level changes from immunoblots performed in (**D**). Mean and standard deviation are shown from at least n = 3 biological replicates. **F)** Immunoblot of the ribosomal protein Rps6 and its phosphorylated form pRps6 were performed in CJY049 background strain, strains deleted for the ArfGEF *GEA1* (Δ*gea1*) or *GEA2* (Δ*gea2*) or Δ*gea2* carrying a *ts*-allele of *GEA1* (*gea1-4 Δgea2*). Pgk1 was used as loading control. Cells were grown to exponential phase at 23°C for 6 h (Expo), shifted for 1 h in media deprived of nitrogen sources (-N), or shifted for 30 or 60 minutes to 37°C. **G)** Relative TORC1 activity based on protein level changes from immunoblots performed in (**F**). Mean and standard deviation are shown from n = 3 biological replicates. Two-way ANOVA with Turkey test was performed. Δ*gea2* Expo *vs* Δ *gea2* 60min *\*\*P:* 0.008037363605601; *gea1-4 Δgea2* Expo *vs gea1-4 Δgea2* 60min \*\*\**P*: 0.000517843389281. **H)** Immunoblot of the ribosomal protein Rps6 and its phosphorylated form pRps6 were performed in RH448 background strain or the *ts*-allele mutant strain *SEC7* (*sec7-1*). Pgk1 was used as loading control. Cells were grown to exponential phase at 23°C for 6 h (Expo), shifted for 1 h in media deprived of nitrogen sources (-N), or shifted for 30 or 60 minutes to 37°C. **I)** Relative TORC1 activity based on protein level changes from immunoblots performed in (**H**). Mean and standard deviation are shown from n = 3 biological replicates. Two-way ANOVA with Turkey test was performed. *sec7-1* Expo *vs sec7-1* 60min \*\*\*\**P*: 0.000000182125903.**J)** Summary of TORC1 activity measured in the different ArfGAP/GEF mutants.

We first determined the TORC1 activity in ArfGAP mutants. Individual deletions of *GLO3* and *GCS1* had no effect on TORC1 activity (**Fig. 2B, 2C**), consistent with our previous finding that *Δglo3* is not rapamycin-sensitive^33^. Glo3 and Gcs1 are partially redundant, and loss of both proteins is lethal. However, a strain carrying only the *ts* allele *GCS1-28* (*Δglo3 Δgcs1* + *GCS1-28*) remains viable. In this strain, TORC1 activity was further reduced following a temperature shift to 37°C (**Fig. 2B, 2C**), indicating that Glo3/Gcs1 function is required for TORC1 activity. The other GAP pair Age1 and Age2 at the TGN seems to regulate TORC1 signaling in the opposite direction as simultaneous deletion of *AGE1* and *AGE2* led to a 50% increase of TORC1 activity after 1h at 37°C (**Fig. 2D, 2E**). Taken together, our data suggest that ArfGAP activity positively regulates TORC1 activity at the Golgi-ER interface, and negatively regulates TORC1 activity at the TGN, suggesting at least two distinct Arf1 functions in the regulation of TORC1. We were wondering whether the decrease of TORC1 activity in Δ*age1* Δ*age2* could be modulated by knocking out *GCS1*. Interestingly, after a shift at 37°C, we could indeed reduce TORC1 activity (**Fig. S2A and B**), indicating that regulation of Arf1 activity at the Golgi is important to balance TORC1 activity.

Next, we measured TORC1 activity in the ArfGEF mutants Δ*gea1*, Δ*gea2* and the double mutant *ts*-strain *gea1-4 Δgea2* (**Fig. 2F**). The individual knock-out strains displayed a slight increase in TORC1 activity at 37°C (**Fig. 2F, 2G**), while in the *ts*-sensitive *gea1-4 Δgea2* double mutant, TORC1 activity was reduced after shift to 37°C. In a loss-of-function *ts*-mutant strain of the ArfGEF Sec7, which acts at the TGN, TORC1 activity was diminished after 60 min incubation at 37°C (**Fig. 2H, 2I**). Thus, TORC1 activity can be potentially regulated at least in two ways, one involving the Golgi-ER shuttle and the one the TGN export pathways. Moreover, our data indicate that cycling between the active and inactive state is necessary for the regulation in the retrograde pathway to the ER, while the second pathways at the TGN might be more reliant on active Arf1 (**Fig. 2J)**.

### Arf1 interacts with TORC1 components in an allele-specific manner

Next, we asked how Arf1 might regulate TORC1. We decided to use an unbiased approach and performed pulldowns from cells grown at either 23°C or after shift to 37°C with either Arf1-GFP or Arf1-11-GFP as a bait (**Fig. 3A**). Components of the TORC1 complex and the TORC1 regulatory complexes SEACAT and SEACIT were enriched in pulldowns of the hyperactive Arf1-11-GFP compared to wild-type Arf1-GFP (**Fig. 3B-F**). This effect was stronger at 37°C compared to 23°C. However, we did not detect components of the EGO complex, which is an activator of TORC1 activity. Besides the expected GO terms such as ‘Transport’, ‘Establishment of localization’ and ‘Localization’, proteins involved in catabolic processes were enriched in the Arf1-11 pulldowns (**Figure 3E, F, Supplementary Table 4**). We have shown previously that autophagy and the UPR are upregulated in *arf1-11*^5^. Our data, however, now suggest a more direct involvement in these processes. To corroborate the interaction between active Arf1 and the TORC1 complex, we performed an immunoprecipitation experiment with the TORC1 subunit Kog1 fused to GFP. Kog1 specifically pulled down the hyperactive mutant Arf1-11, indicating that the TORC1 complex only interacts with the activated form of Arf1 (**Fig. 3G**).

**Figure 3.**
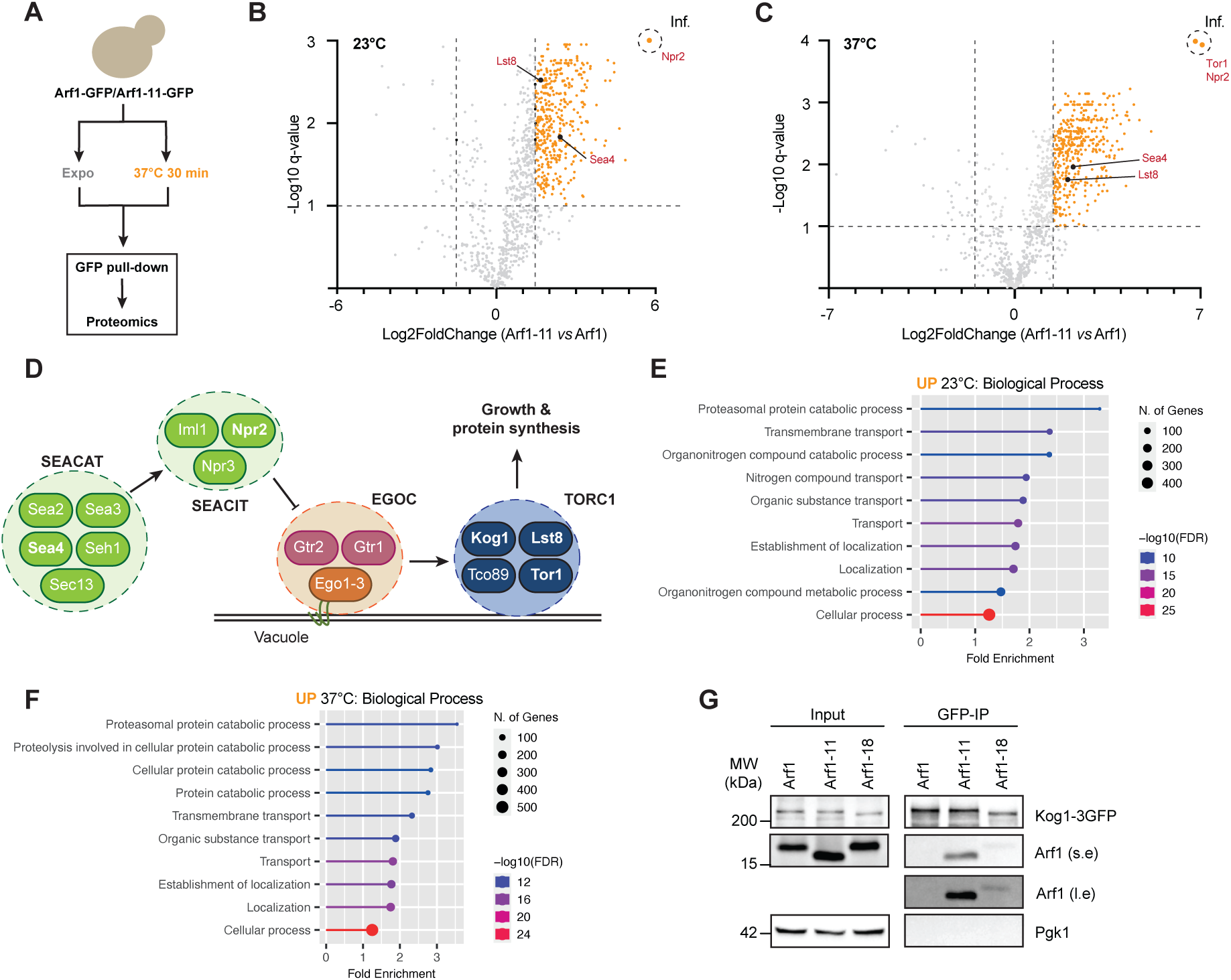
Arf1 interacts with TORC1 components in an allele-specific manner. **A)** Schematic depiction of the strategy used to identify pathways deregulated in the *arf1-11 ts*-allele mutant strain by proteomics using Arf1 and Arf1-11 fused to GFP. **B-C)** Volcano plot obtained from Arf1- and Arf1-11-GFP strains grown to exponential phase at 23°C (**B**) or shifted to 37°C for 30 minutes (**C**). All increased pulled-down proteins in Arf1-11-GFP are shown in orange. Upregulated proteins related to TORC1 are shown. See also Supplementary Table 2 and 3. **D)** Schematic depiction of TORC1 complex regulation by EGOC, SEACAT and SEACIT complexes in yeast. **E-F)** Gene Ontology terms associated with yeast biological processes based on increased proteins by pull-down at 23°C (**E**) and 37°C (**F**). **G)** Pull-down assay performed on the TORC1 kinase Kog1 fused to 3xGFP. Immunoblot of Kog1-3GFP in the WT Arf1 strain and related alleles was performed. Pgk1 was used as control.

### *arf1* mutants induce foci containing Kog1

Since Arf1 interacts with TORC1 in an activity-dependent manner, we tested next whether Arf1 mutants would also affect TORC1 localization. Therefore, we tagged the TORC1 component Kog1 with 3mCherry in strains in which Arf1 or Arf1 mutants were tagged with GFP (**Fig. 4A**). At permissive temperature, Kog1 localization was mostly found at the vacuole, where active TORC1 resides. In addition, we also observed Kog1 in discrete foci distinct from the vacuole in the hyperactive Arf1 strains (**Fig. 4A**; arrows). After shift to the restrictive temperature, such Kog1 foci were observed in all three strains by widefield microscopy but more noticeable by high-resolution microscopy (**Fig. 4A and B**). The abundance of these foci was more pronounced in the mutants compared to wild type (**Fig. 4C**), consistent with a greater reduction of TORC1 activity after temperature shift (**Fig. 1D and E**). Moreover, they were also observed in cells deprived of nitrogen (-N) at 23°C and after shift 30 min at 37°C, or in the absence of glucose (-D). However pharmacological inhibition of TORC1 activity with rapamycin did not dissociate TORC1 from the vacuole (**Fig. S3A**). Taken together, these structures are reminiscent of TORC1 organized in inhibited domains^18^ (TOROIDS) and TORC1 bodies^19^.

**Figure 4.**
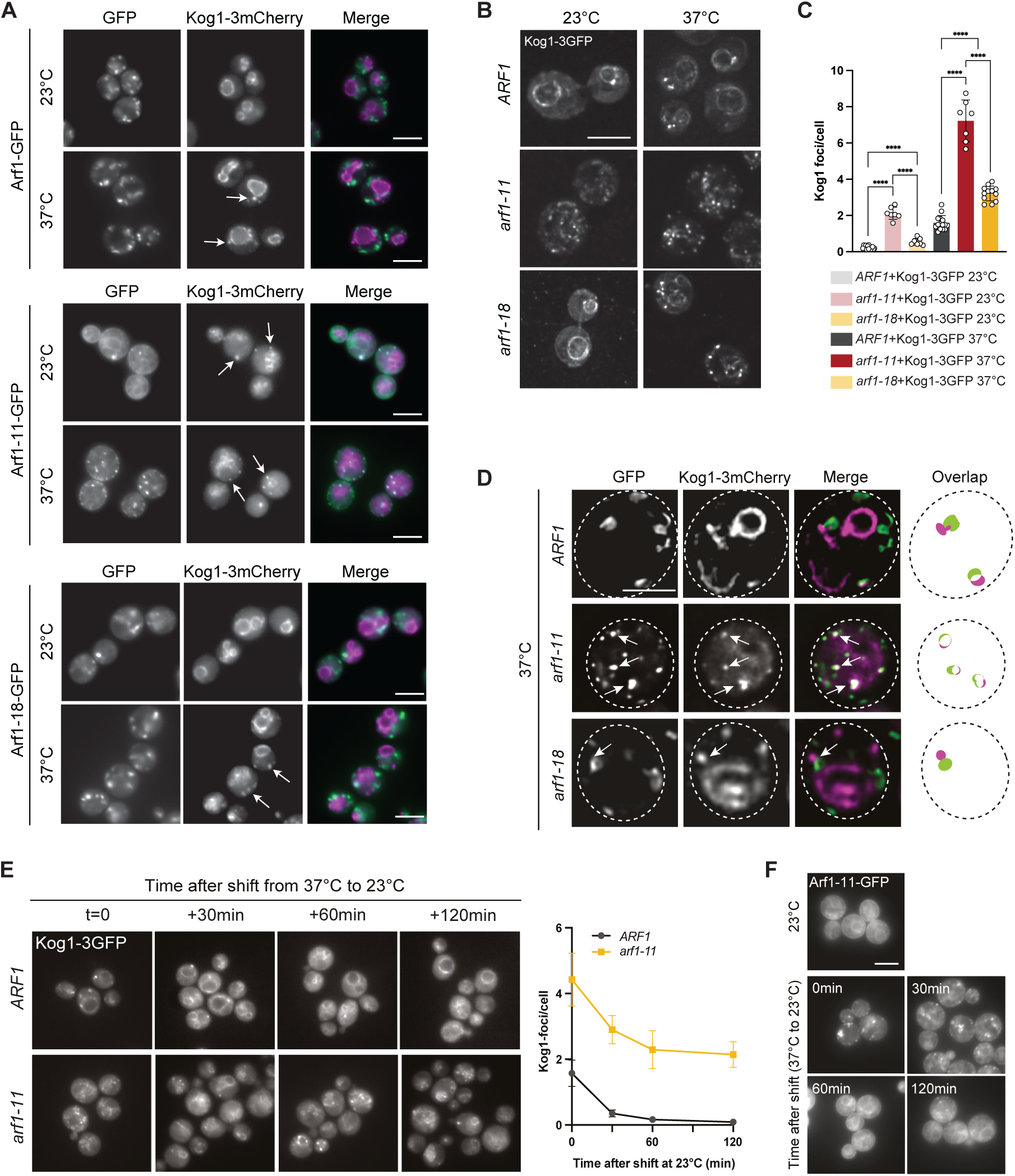
*arf1* mutants induce foci containing Kog1. **A)** Widefield microscopy of Arf1-, Arf1-11- and Arf1-18-GFP strains expressing Kog1 fused to 3xmCherry and grown to exponential phase at 23°C for 6 h or shifted for 30 minutes to 37°C. Arrows indicate Kog1 foci. Scale bar: 5 µm. **B)** FEI-MORE high-resolution widefield microscopy of *ARF1, arf1-11* and *arf1-18 ts*-allele strains expressing Kog1 fused to 3xGFP and grown to exponential phase at 23°C for 6 h or shifted for 30 minutes to 37°C. Scale bar: 5 µm. **C)**Measurements of Kog1-3GFP foci number per cell in all three strains at both temperatures tested in (**B**). Mean and standard deviation are shown from at least n = 3 biological replicates. Unpaired *t*-test. *ARF1* 23°C vs *arf1-11* 23°C \*\*\*\**P*: 0,000000000000026; *ARF1* 23°C vs *arf1-18* 23°C \*\*\*\**P*: 0,000050220723566; *arf1-11* 23°C vs *arf1-18* 23°C \*\*\*\**P*: 0,000000147803041; *ARF1* 37°C vs *arf1-11* 37°C \*\*\*\**P*: 0,000000000000192; *ARF1* 37°C vs *arf1-18* 37°C \*\*\*\**P*: 0,000000000188712; *arf1-11* 37°C vs *arf1-18* 37°C \*\*\*\**P*: 0,000000003381370. **D)** FEI-MORE high-resolution widefield microscopy of Arf1-, Arf1-11-and Arf1-18-GFP strains expressing Kog1 fused to 3xmCherry and grown to exponential phase at 23°C and shifted for 30 minutes to 37°C. Arrows indicate Kog1 foci colocalizing or juxtaposed to Arf vesicles. Scale bar: 5 µm. **E)** Widefield microscopy of ARF1 and *arf1-11* strains expressing Kog1-3GFP, grown to exponential phase at 23°C for 6 h, then shifted for 30 minutes to 37°C, and shifted back to 23°C for the indicated time points. The number of Kog1 foci were measured for each time point. Mean and standard deviation are shown from at least n = 3 biological replicates. **F)** Widefield microscopy of Arf1-11-GFP grown in the same conditions as (**E**).

Our data appeared to suggest that at least a part of the Kog1 structures also contained Arf1 (**Fig. 4A**). To further explore this possibility, we imaged our strains by high-resolution microscopy at the restrictive temperature. As previously reported, Arf1 and Arf1-18 are mostly present on the Golgi, while Arf-11 localizes predominantly to lipid droplets (LD) at 37°C^10^. In the WT strain and in *arf1-18*, Kog1 was either juxtaposed or partially overlapping with Arf1 (**Fig. 4D**). In contrast, in *arf1-11* cells, almost all Kog1 structures were colocalizing with Arf1-11 (**Fig. 4D**), suggesting that active Arf1 could be recruited to Kog1 foci. We, thus, hypothesized that Kog1 foci contain inactive TORC1 and other components and could potentially act as a buffer. If so, alleviating the stress, i.e. shifting to the permissive temperature, should reduce the number of these foci. We indeed observed that the kinetics of disappearance were similar in wild-type and *arf1-11* cells (**Fig. 4E**). *arf1-11* cells retained more Kog1 foci because TORC1 activity is already affected at 23°C in this mutant. Interestingly, the dissolution kinetics of the Kog1 foci was very similar to the time needed for most of the Arf1-11 signal to relocalize from LDs to the ER^10^ (**Fig. 4F**). Taken together, our data imply that dysregulation of Arf1 activity, which in turn inactivates TORC1 leads to the formation of Kog1 structures.

### Kog1 foci represent a distinct reservoir for TORC1

Next, we aimed to determine whether *arf1-11* induces either TOROIDS or TORC1 bodies^19,34^. Both structures sequester inactive TORC1, including the Tor1 kinase. Therefore, we tested whether Tor1 and Kog1 colocalize in the Kog1-3GFP foci. In the *ARF1* strain, GFP-Tor1 localized around the vacuolar rim as expected in cells grown at 23°C and 37°C (**Fig. S4A**) and additionally in foci reminiscent of the Kog1 foci at 37°C. (**Fig. S4A,** arrows). We found that 50% of cells observed harbored foci containing both Kog1 and Tor1 (**Fig. 5A and B**). TOROIDs are composed of an oligomer of TORC1 complex units that form a barrel shaped structure^18,34^, while the precise composition of TORC1 bodies is less clear^19,20^. It is also possible that TORC1 bodies and TOROIDs refer to the same structure. Still, the *arf1-11* strain induced Kog1 foci containing Tor1 did not appear to be barrel-shaped at microscopic resolution. TORC1 bodies assemble on the vacuolar rim and require the EGO complex for their assembly with which they co-localize^19,20^. To test whether the Kog1 foci are TORC1 bodies, we determined the localization of the EGOC component Gtr1 after shift to 37°C for 30 min. We did not see any Gtr1 structures colocalizing with Kog1-3GFP dots in the cytoplasm in *ARF1*, *arf1-11* and *arf1-18* strains (**Fig. 5C**), suggesting that the Kog1 foci are distinct from TORC1 bodies. We next asked whether upstream components of the TORC1 regulatory module might be co-localizing with the Kog1 foci. GATOR1 and GATOR2 homologues in yeast, namely SEACAT and SEACIT regulate Gtr1 and the Ego complex^35,36^. The SEACAT subunit Sea4 fused to mCherry did not form structures like Kog1/Tor1-3GFP, and the fluorescent signal stayed on the vacuolar ring in *ARF1*, *arf1-11* and *arf1-18* strains (**Fig. 5D**). Similarly, the SEACIT subunit Iml1-3mCherry did not form any foci (**Fig. S4B**). In contrast, the SEACIT component Npr2 formed similar structures as Kog1/Tor1 in all three *ARF1* strains after 30 min shift at 37°C (**Fig. 5E**). Interestingly, Npr2-3mCherry and Kog1-3GFP colocalized in the *ARF1* and *arf1-18* strains, but not in the *arf1-11* mutant allele where they were juxtaposed but never overlapped, almost as if the presence of Arf1-11 and Npr2 in the Kog1 foci was mutually exclusive. Our data suggest that the Kog1 foci are distinct from TORC1 bodies and TOROIDs in terms of composition and/or assembly pathway. Therefore, we decided to name them ART (*Ar*f1-dependent *T*ORC1) bodies.

**Figure 5.**
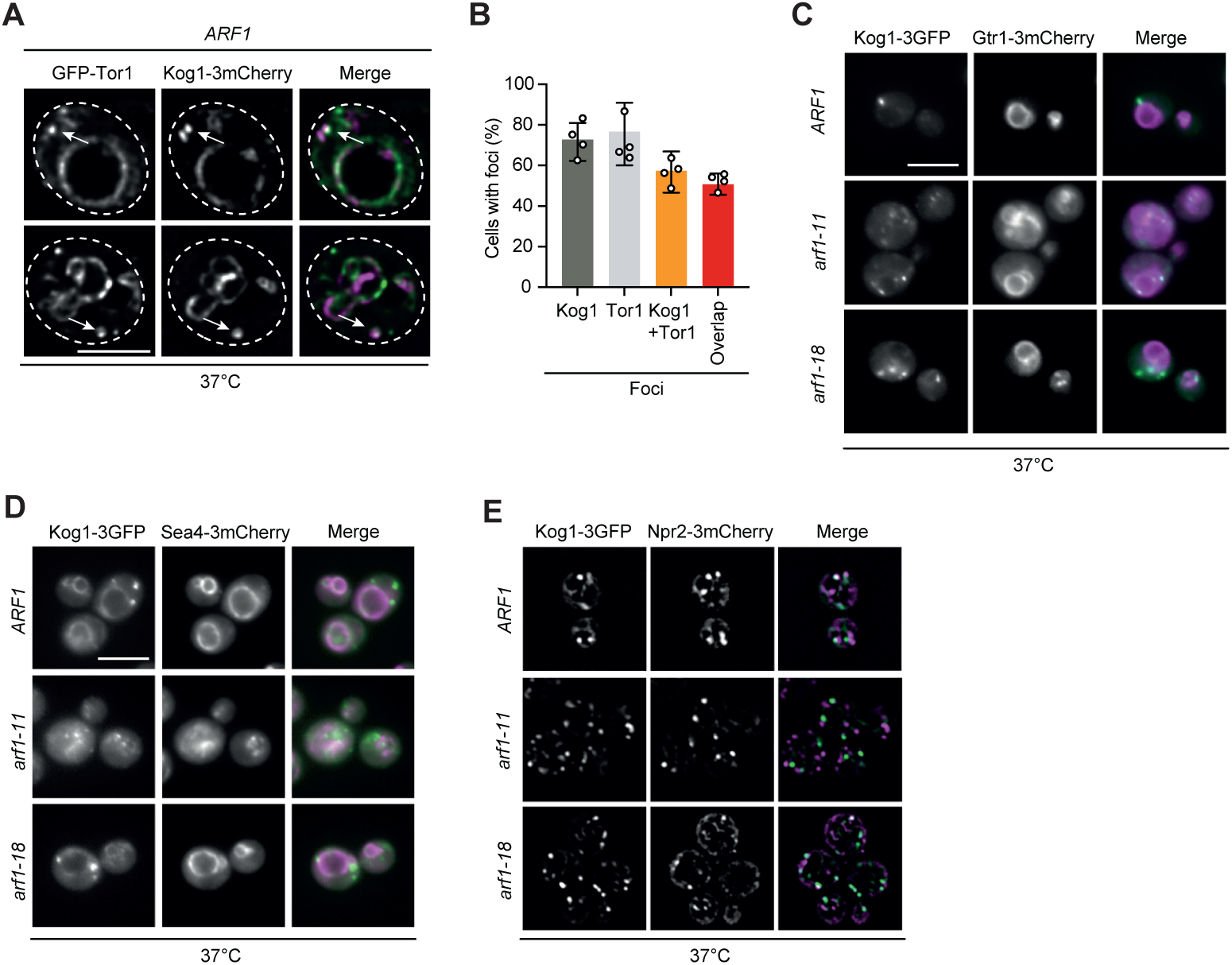
Kog1 foci represent a distinct reservoir for TORC1. **A)** FEI-MORE high-resolution widefield microscopy of *ARF1* strain expressing GFP-Tor1 and Kog1-3mCherry grown to exponential phase at 23°C and shifted for 30 minutes to 37°C. Arrows indicate Kog1 foci colocalizing with Tor1. Scale bar: 5 µm. **B)** Measurements of cells containing either Kog1 or Tor1 foci alone, or both, and how much of them colocalize. Mean and standard deviation are shown from n = 4 biological replicates. **C)** Widefield microscopy of *ARF1, arf1-11* and *arf1-18 ts*-allele strains expressing Kog1-3GFP and Gtr1-3mCherry grown to exponential phase at 23°C for 6 h and shifted for 30 minutes to 37°C. Scale bar: 5 µm. **D)** Widefield microscopy of *ARF1, arf1-11* and *arf1-18 ts*-allele strains expressing Kog1-3GFP and Sea4-3mCherry grown to exponential phase at 23°C for 6 h and shifted for 30 minutes. Scale bar: 5 µm. **E)** Widefield microscopy of *ARF1, arf1-11* and *arf1-18 ts*-allele strains expressing Kog1-3GFP and Npr2-3mCherry grown to exponential phase at 23°C for 6 h and shifted for 30 minutes to 37°C. Images were processed with Gaussian blur in Fiji. Scale bar: 5 µm.

Besides its localization on the vacuole where it controls protein synthesis and cell growth^37^, TORC1 has been recently shown to be routed to signaling endosomes where it regulates micro- and macro-autophagy^38–40^. Considering that wild-type cells contain on average 1.5 ARTs per cell (**Fig. 4C**), we asked whether ARTs were signaling endosomes. Ivy1 is a marker for these signaling endosomes induced under nutrient starvation^38^. Nevertheless, we never observed co-localization of Ivy1-3mCherry with Kog1-3GFP, apart from the vacuole, after temperature shift (**Fig. S5A**), suggesting that ARTs are not signaling endosomes. Kog1-GFP has also been shown to be stored in stress granules upon heat stress at 46°C for 30 min^41^. We thus tested whether ARTs could be stress granules or processing bodies (PBs) using Pab1 and Dcp1, respectively, as markers. Although *arf1-11* and *arf1-18* strains generated PBs at 23°C and 37°C^5^, none of them colocalized with Kog1-3GFP (**Fig. S5B**). Another possibility is that Kog1-3GFP is sequestered in stress granules (SGs), as we induce a slight heat stress by shifting cells to 37°C^41^. However, Pab1-mScarlet did not form SGs at 37°C in any of the strain tested (**Fig. S5C**) but did after 30 min incubation at 46°C to which Kog1-3GFP rarely localized (**Fig. S5D**). Moreover, ARTs did not disassemble after 1,6-hexanediol treatment, indicating they do not represent dynamic condensates (**Fig. S5E**). We conclude that ARTs are not signaling endosomes, PBs or SGs but represent structures that sequester TORC1 and at least one other component of the SEACIT complex.

### ER-anchored Arf1-11 partially rescues TORC1 activity at the restrictive temperature

Our data above suggest that Arf1 may regulate TORC1 activity in two independent ways: one by regulating the transport from the TGN to the vacuole and second through the ER-Golgi shuttle (**Fig. 2**). ARTs were not confined to vacuolar proximity but in contrast appeared to be mobile. We wondered whether ARTs might contact the ER and found that 70-80% of ARTs were juxtaposed to the ER in wild-type and mutant strains (**Fig. 6A-C**). Average contact times between ARTs and the ER were significantly longer in *arf1-11* compared to *ARF1* and *arf1-18* strains, with several contacts lasting at least two minutes, which was the total duration of the movies (**Fig. 6D**). Arf1-11 is enriched on LDs at 37 °C and since TORC1 has been shown to regulate LDs formation and triacylglycerol levels^42^, we asked if Kog1-3GFP also colocalized with LDs. However, we could not see any overlap between Kog1 and the LD marker Erg6 (**Fig. S6A**). Our data suggest that Arf1 on the ER might influence TORC1 activity. We previously described that artificially anchoring Arf1-11 on the ER at the non-permissive temperature, restores fatty acids transport and mitochondrial homeostasis^10^ (**Fig. 6E**). We thus measured TORC1 activity in strains where either Arf1 or Arf1-11 was anchored to the ER in a Δ*arf1* background. ER-anchored Arf1-11 partially rescued the loss of TORC activity of the *arf1-11* mutant at 37°C (**Fig. 1D, 1E, Fig. 6F**). Thus, ER-localized Arf1 might positively regulate TORC1 activity.

**Figure 6.**
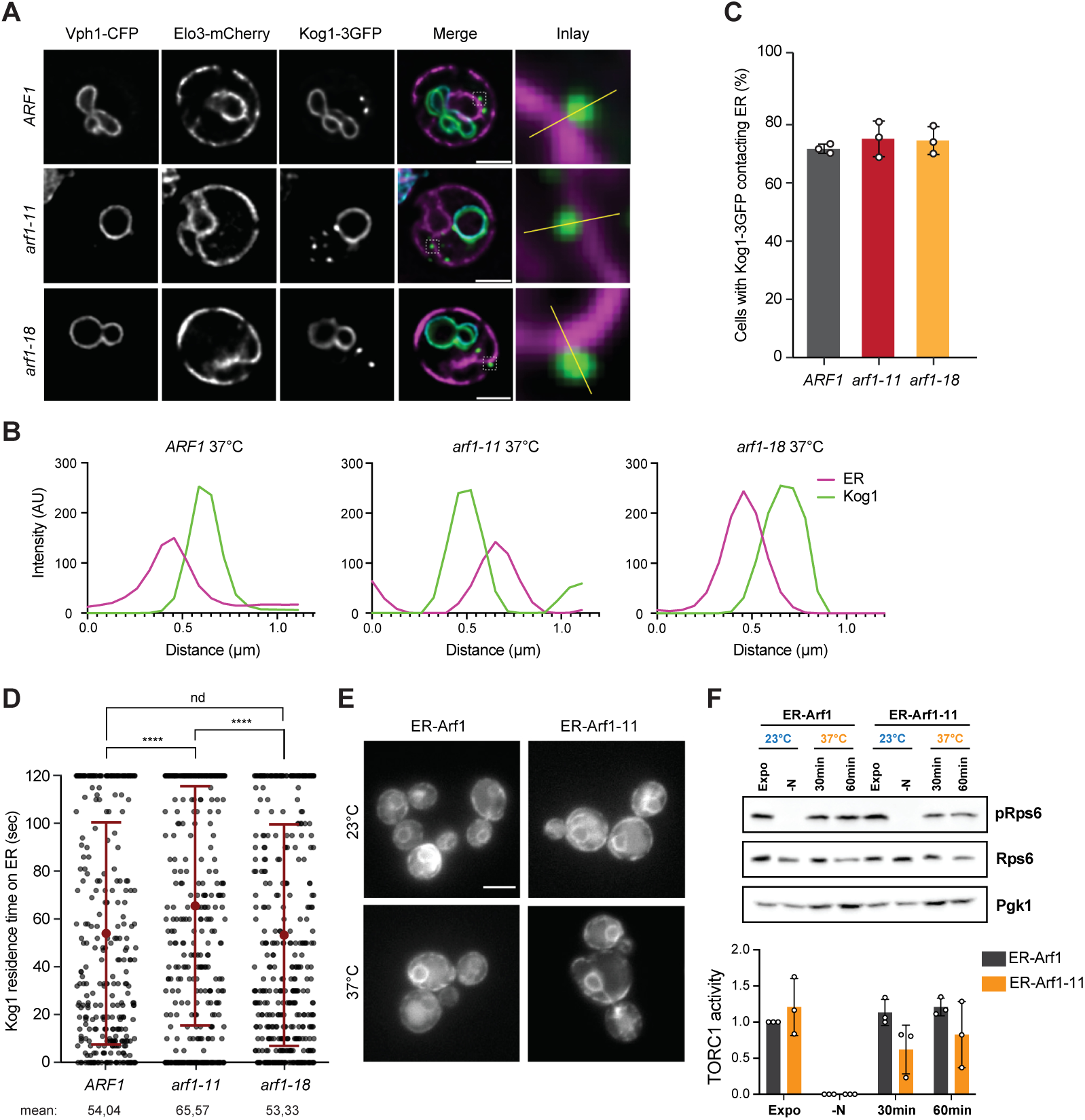
ER-anchored Arf1-11 partially rescues TORC1 activity at the restrictive temperature. **A)** FEI-MORE high-resolution widefield microscopy of *ARF1, arf1-11* and *arf1-18 ts*-allele strains expressing Kog1-3GFP together with the vacuolar marker Vph1-CFP and the ER marker Elo3-mCherry grown to exponential phase at 23°C and shifted for 30 minutes to 37°C. Scale bar: 5 µm. **B)** Line plots from the individual channels from the inlays in (**A**) encompassing the ER and Kog1 foci. Intensities are shown as arbitrary units (a.u.). **C)** Measurements of *ARF1, arf1-11* and *arf1-18* cells with Kog1-3GFP foci in contact with the ER. Mean and standard deviation are shown from n = 3 biological replicates. **D)** Kog1 foci residence time on the ER measured from 120 minutes long movies from cells grown to exponential phase at 23°C and shifted for 30 minutes to 37°C. Individual dots represent individual Kog1 foci analyzed. Mean (red dot) and standard deviation are shown from n = 3 biological replicates. Welch two samples *t*-test with Bonferroni correction was performed. *ARF1 vs arf1-11* \*\*\*\**P*: 0.0009793; *arf1-11 vs arf1-18* \*\*\*\**P*: 0.0001677. nd: not different. **E)** Widefield microscopy images from Δ*arf1* strain expressing ER-anchored Arf1- or Arf1-11-GFP in cells grown to exponential phase at 23°C or shifted for 30 minutes to 37°C. Scale bar: 5 µm. **F)** Immunoblot of Rps6 and its phosphorylated form pRps6 performed in ER-Arf1-GFP and ER-Arf1-11-GFP strains. Pgk1 was used as loading control. Cells were grown to exponential phase at 23°C for 6 h (Expo), shifted for 1 h in media deprived of nitrogen sources (-N), or shifted for 30 or 60 minutes to 37°C. Mean and standard deviation are shown from n = 3 biological replicates.

### Kog1 function is regulated by Arf1 alleles

To gain more insights about how Arf1 regulates TORC1 at the ER, we decided to measure TORC1 hysteresis, which is the capacity of TORC1 to be reactivated after stress-dependent inhibition^19^. We measured TORC1 re-activation after 5 and 15 min of nutrients addition after a 1h inactivation by nitrogen starvation (-N) at 23°C (**Fig. 7A, B**). Under these conditions, TORC1 activity was 3-fold increased in the ER-localized *arf1-11* hyperactive mutant over Golgi-localized wild-type Arf1. In contrast, TORC1 was not re-activated in the loss-of-function mutant *arf1-18* (**Fig. 7A, 7B**). Arf1-18 still localized to the Golgi under these conditions (**Fig. S1G**). We have previously shown that Glo3 can be phosphorylated by Snf1 kinase, which indirectly regulates TORC1 activity^43^. Moreover, the non-phosphorylatable Glo3 destabilized and the phosphomimetic Glo3 stabilized Arf1 on membranes^43^. We therefore tested whether Glo3 phosphomutants would affect TORC1 re-activation. Expression of both the non-phosphorylatable and phosphomimetic mutants of Glo3 led to a reduction of TORC1 activity already in the absence of stress, which was slightly more pronounced in the case of the phosphomimetic mutant (**Fig. 7C, D**). This trend persisted after reactivation, indicating that both phosphorylated and non-phosporylated forms of Glo3 are required to allow restoration of TORC1 activity after stress. Taken together, our data indicate that active, ER-localized Arf1 is beneficial for the re-establishment of TORC1 activity after stress.

**Figure 7.**
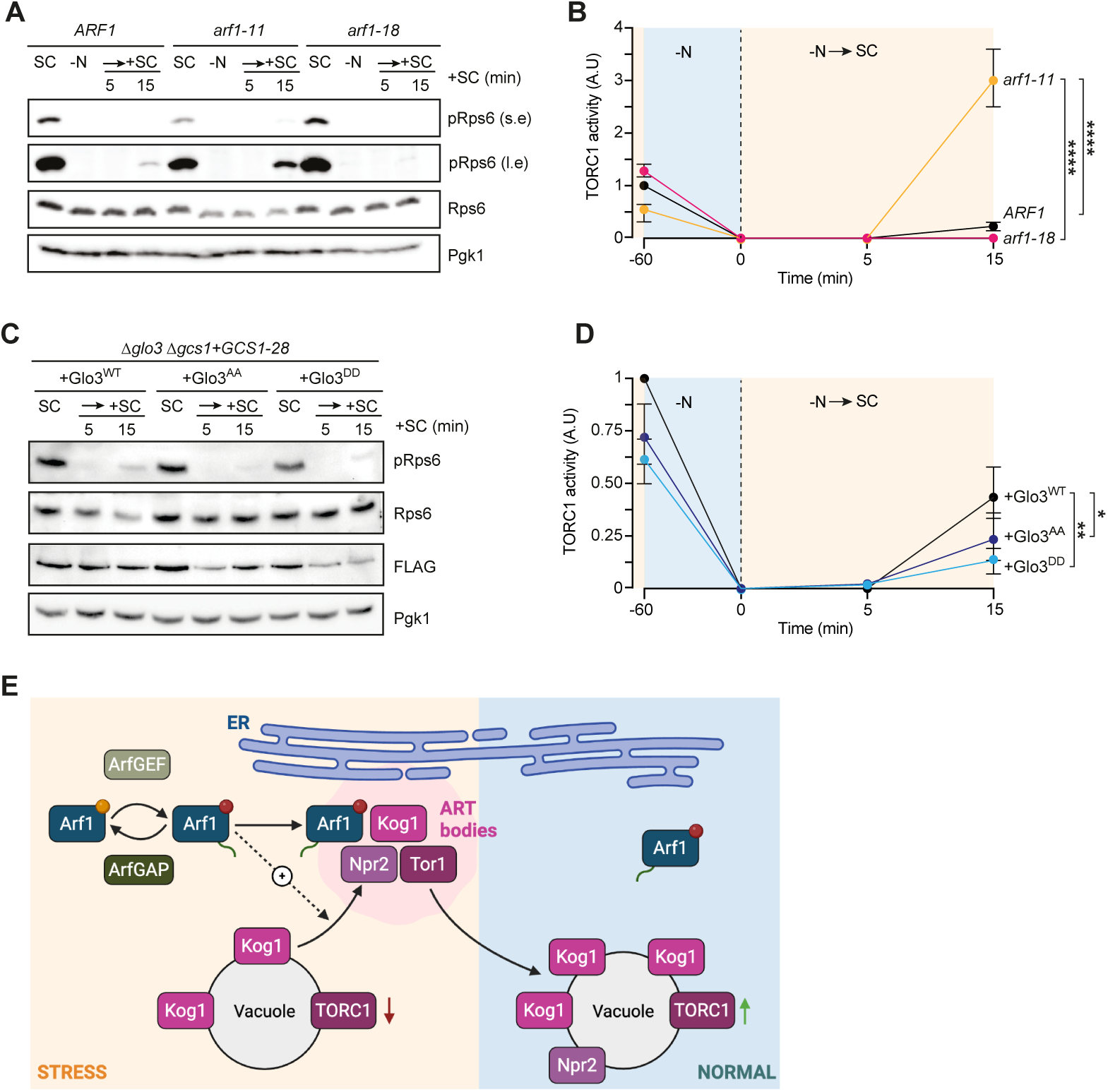
Kog1 function is regulated by Arf1 alleles. **A)** Immunoblot of Rps6 and its phosphorylated form pRps6 performed in *ARF1, arf1-11* and *arf1-18* strains. Pgk1 was used as loading control. Cells were grown to exponential phase at 23°C for 6 h in synthetic media (SC), shifted for 1 h into media deprived of nitrogen sources (-N) or shifted into -N for 1 h and then shifted back to SC media at 23°C for indicated times. **B)** TORC1 activity was measured and plotted based on experiments performed in (**A**). Mean and standard deviation are shown from n = 3 biological replicates. Two-way ANOVA with Turkey test. *ARF1 vs arf1-11* +SC \*\*\*\**P*: 0.000000249591756 and *arf1-11 vs arf1-18* +SC \*\*\*\**P*: 0.000000139756537. s.e: short exposure; l.e: long exposure. **C)** Immunoblot of Rps6 and its phosphorylated form pRps6 performed in Δ*glo3 Δgcs1+GCS1-28* strain expressing WT Glo3, non-phosphorylatable Glo3 (Glo3^AA^) and phosphomimetic Glo3 (Glo3^DD^) fused with a FLAG tag from a plasmid. Pgk1 was used as loading control, and anti-FLAG to assess Glo3 expression. Cells were grown to exponential phase at 23°C for 6 h in synthetic media (SC), shifted for 1 h into media deprived of nitrogen sources (-N) or shifted into -N for 1 h and then shifted back to SC media at 23°C for indicated times. **D)** TORC1 activity measured and plotted based on experiments done in (**C**). Mean and standard deviation are shown from n = 3 biological replicates. Two-way ANOVA with Turkey test. Δ*glo3 Δgcs1+GCS1-28* + Glo3 WT *vs* Δ*glo3 Δgcs1+GCS1-28* + Glo3^AA^ \**P*: 0.0449; Δ*glo3 Δgcs1+GCS1-28* + Glo3 WT *vs* Δ*glo3 Δgcs1+GCS1-28* + Glo3^DD^ \*\**P*: 0.0084. s.e: short exposure; l.e: long exposure. **E)** Model of the mechanism on how Arf1 regulates TORC1 activity under normal and stress conditions. Figure was made with BioRender.com. For more information, see discussion section.

## Discussion

Our study reveals a pivotal role for Arf1 in modulating the formation of distinct cytoplasmic bodies, which sequester TORC1 components, when TORC1 activity is dampened. Moreover, we establish that Arf1 activity influences TORC1 hysteresis. We provide evidence that in yeast, Arf1 colocalizes with TORC1 components Kog1 and Tor1 following mild heat stress (37°C for 30 min) and in the presence of a hyperactive Arf1 mutant. This colocalization extends beyond the vacuolar membrane, occurring in discrete cytoplasmic bodies near the ER. Interestingly, while steady-state Arf1-Kog1/Raptor interactions are barely detectable, we show that a hyperactive mutant of Arf1 stabilizes this interaction. An Arf1 loss-of-function mutant also exhibits mild stabilization - suggesting that proper GTP-GDP cycling is critical for TORC1 spatial dynamics.

We observed assemblies containing Kog1 and Tor1, as well as Arf-11. Considering that they are structurally and functionally distinct from heat-induced stress granules^41^, signaling endosomes^40^, and TOROIDs^18^, we decided to name these foci Arf1-related TORC1 bodies (ART-bodies or ARTs for short). We propose that TORC1 components can be sequestered in different ways and in structures of distinct composition in a context-dependent manner upon TORC1 inactivation. Our findings suggest that ART-bodies formation correlates with reduced TORC1 activity and increased hysteresis, as observed in *arf1-11* versus *arf1-18* strains, as well as in the Glo3^S289–298^ phospho-mutant. Npr2 did not localize to ARTs in the *arf1-11* strain. We hypothesize that activated Arf1 is involved in the selective recruitment of proteins to ART to keep TORC1 inactive. Altogether, our data suggest that Arf1 is involved in the formation of ARTs under stress conditions, regulates protein composition of ARTs to keep Tor1 inactive and to prevent TORC1 premature reactivation after stress released (**Fig. 7E**). In this scenario, Arf1 function is dependent on its GTPase cycle and is tightly regulated by ArfGAPs and ArfGEFs.

Emerging evidence suggest that the spatial segregation of mTORC1 activity is essential for distinct cellular functions, with phosphorylation targets varying by subcellular localization^44,45^. Although the mechanisms by which Arf1 regulates ARTs formation remain unclear, we could show that this process is independent of the yeast Rag-GTPase Gtr1/Gtr2, similar to what has been described for Arf1 regulation on TOR activity^46^. Given that there are at least three distinct molecular assemblies for inactive Tor1, it is likely that these assemblies are not strictly storage spaces but might also have individual functions. What these functions are, remains to be determined.

Our data indicate that Arf1-mediated regulation of TORC1 activity is at least twofold: First, Arf1 acts are the TGN in delivery of vacuolar proteins, essential for vacuolar homeostasis. Second, Arf1 acts at the Golgi-ER-vacuole interface. In this latter case, cycling between GTP and GDP bound forms are essential. Hence, Arf1 act as a hub to regulate TORC1 activity, integrating proper trafficking towards vacuole and nutrient sensing at the ER-vacuole interface.

Our study provides new mechanistic insights into the spatial regulation of TORC1 in yeast, highlighting how Arf1 orchestrates TORC1 localization and activity in response to metabolic and stress cues. These findings have broad implications for understanding how intracellular signaling pathways integrate environmental stimuli, and how small GTPases coordinate the dynamics of multimeric complexes such as TORC1. Future investigations into the molecular determinants of TORC1 sequestration and reactivation will be instrumental in elucidating the broader principles of spatial signal transduction in eukaryotic cells.

## Materials and Methods

### Strains, media and plasmids

Yeast strains were either grown in rich media composed of 1% w/v yeast extract, 1% (w/v) peptone, 40 mg L^−1^ adenine, 2% (w/v) glucose (YPD) or 2% glycerol (YPGly), or in synthetic complete medium (HC) composed of 0.17% (w/v) yeast nitrogen base with ammonium sulfate and without amino acids, 2% (w/v) glucose and mixtures of amino acids (MP Biomedicals) depending on the auxotrophies used for selection. Unless otherwise indicated, cells were grown at 23°C and a subset was shifted to 37°C for 30 min before analysis. YPD plates with rapamycin contained 5 ng/mL rapamycin (SelleckChem) and 2% (w/v) agar. Media deprived of glucose (-D) is composed of 1% w/v yeast extract, 1% (w/v) peptone and 40 mg L^−1^ adenine (YP).

### Yeast transformation

Three units of OD600 of yeast cells were grown in appropriate YPD or HC media to mid-log phase. Cells were spun down and washed in 1 volume of 1× TE and 10 mM LiAc. The pellet was then resuspended in 350 µL of transformation mix (1× TE, 100 mM LiAc, 8.5% (v/v) single-stranded DNA and 70% (v/v) PEG3000), incubated with DNA (PCR product or plasmid) for 1 h at 42 °C, spun down (30 s at 10,000*g* at RT) and resuspended in 100 μL of YPD or HC media, and cells were plated onto selective media and incubated at 23°C or 30°C. Genomic tagging was performed according to standard procedures^47^.

### Microscopy

Fluorescence and DIC images were acquired with an ORCA-Flash 4.0 camera (Hamamatsu) mounted on an Axio Imager.M2 fluorescence microscope with a 63× Plan-Apochromat objective (Carl Zeiss) and an HXP 120C light source with ZEN 2.6 software. High-resolution images were acquired with an ORCA-Flash 4.0 camera (Hamamatsu) mounted on a FEI-MORE microscope with a 100× U Plan-S-Apochromat objective (Olympus). Image processing was performed and analyzed with Fiji software. Measurements of ART-bodies number and duration of ER contacts were perfomed with Fiji software.

### TORC1 activity and hysteresis assays

To assess TORC1 activity and inhibition by immunoblot, cells were grown overnight in YPD to reach mid-log phase, diluted to 0.2 OD600 and grown for 6 h in fresh YPD. Cells were harvested and washed twice in nitrogen starvation media (-N; 1,7g/L YNB w/o amino acids and ammonium sulfate (MP Biomedicals) and 2% (w/v) glucose) and incubated in equal volumes of -N media for 1 h at 23°C or 37°C. For starvation/refeed experiments, cells were grown in YPD media, washed twice in -N media and grown in -N starvation media for 1 h at 23°C or 37°C, spun 2 min at 2,100 xg at RT and resuspended in equal volumes of HC media supplemented with 2% glucose for the indicated time points.

### ART-bodies induction and chemical treatments

To test for the induction of ART-bodies in various conditions, cells were grown overnight in YPD to reach mid-log phase, diluted to 0.2 OD600 and grown for 6 h in fresh HC media. Then cells were washed twice in appropriate media and grown for 1 h in -N starvation media at 23°C or shifted for an additional 30 min at 37°C, or in YPD and in the presence of 5 ng/mL of rapamycin for 1h at 23°C or with an additional shift at 37°C for 30 min, or in YP media deprived of glucose (-D) for 15 min. To test the LLPS properties of ART-bodies, yeast cells were grown overnight in HC media to reach mid-log phase, diluted to 0.2 OD600 and grown for 6 h in fresh HC media. Then 1M 1,6-hexanediol (Sigma-Aldrich) was added to the cultures for the indicated time-points.

### Protein extraction and immunoblot analysis

Ten mL of mid-log grown cultures were lysed using the TCA-NaOH method. Briefly, cells were spun 2 min at 2,100 xg at RT, and the pellet resuspended in 500 µL of ddH2O. Then 50 µL of NaOH (1.85 M) were added and the mixture was incubated for 10 min on ice. Then 50 µL of 100% TCA was added and the mixture was incubated for 10 min on ice. Precipitates were then centrifuged 10 min at 13,000 xg at 4°C. Supernatants were removed and pellets resuspended in 2x Laemmli buffer. Equal protein concentration were loaded on 10%, 12% or 15% SDS–PAGE and transferred onto 0.45 µm nitrocellulose membranes (Amersham). Membranes were blocked with TBST (20 mM Tris, 150 mM NaCl, pH 7.6 and 0.1% Tween20) with 5% non-fat dry milk for 30 min and incubated with anti-Rps6 primary antibody (1:5,000,Gift from Loewith lab raised against the RVFFDKRIGQEVDGE peptide in guinea-pig), anti-pRps6 ser235-236 (1:10,000, Cell Signaling #2211) or anti-Pgk1 primary antibody (1:5,000, Invitrogen clone 22C5D8) overnight at 4 °C, followed by 2 h incubation with horseradish peroxidase (HRP)-conjugated secondary antibodies (1:10,000 anti-mouse, Invitrogen 31430; 1:10,000 anti-rabbit, Sigma-Aldrich A0545; 1:10,000 anti-goat,) in TBST. Chemiluminescence signals were detected using Immobilon Western HRP Substrate (Millipore) and imaged using a FusionFX (Vilber Lourmat).

### Arf1 immunoprecipitation

Yeast cultures of 3×10^5^ cells were grown overnight in 100 mL of YPD media at 23°C. When cultures reached mid-log phase, cells were split in half to be processed for IP or shifted 30 min at 37°C. Cells were spun down, and lysed in 300 µL of lysis buffer (25 mM Tris-HCl pH7.5, 150 mM NaCl, 2 mM EDTA, 0.6% Triton ×100, 1 mM DTT, 1x Halt protease inhibitor cocktail (ThermoScientific)) in the presence of 100 µL of glass beads (0.25–0.5 mm; ROTH) and lysed at 4°C for 20 min on a table-top vortexer. Unbroken cells and debris were pelleted 5min at 3,000 xg at 4°C, and the supernatant incubated with pre-equilibrated 25 µL anti-GFP magnetic beads (Chromotek) for 1h at 4°C on a rotating stand. Beads were then washed 4 times with wash buffer (25 mM Tris-HCl pH7.5, 150 mM NaCl, 2 mM EDTA, 0.06% Triton X100, 1 mM DTT, 1x Halt protease inhibitor cocktail) and elution was performed by adding 25 µL of Laemmli w/o Coomassie blue (125 mM Tris-HCl pH 7.5, 5% SDS and 10% β-mercaptoethanol) to the beads and incubating 5 min at 95°C.

### Co-immunoprecipitation

For Kog1 co-immunoprecipitation, 250 mL of mid-log grown yeast cells cultures were spun down and cells resuspended in 4 mL of breaking buffer (1x PBS, 0.5% (v/v) Tween20, 10% (w/v) glycerol, 2 mM EDTA, 5 mM MgCl2, 2x Halt protease inhibitor cocktail (ThermoScientific), 0.1 mM PMSF (100 mM stock)). Cells were then transferred to 15 mL Falcon tubes supplemented with 1 mL glass beads (0.25–0.5 mm; ROTH) and lysed at 4 °C in with a Fastprep (MP Biomedicals), 8 times 30 sec with 1 min resting time in between each run. Cell debris and unbroken cells were pelleted by centrifugation 3,000 xg for 5 min at 4 °C. The supernatant was then transferred into a new 15 mL incubated with 1 mM DSP for 30 min at 4°C on a rocking bed. Crosslinking was then quenched by adding 100 mM Tris-HCl pH 7.5 for 30 min at 4°C on a rocking bed. Cell extracts were supplemented with 25 µL anti-GFP magnetic beads (Chromotek) and incubated O/N at 4 °C under rotation. Beads were then washed four times in lysis buffer, and proteins were eluted with 50 µL of 2x Laemmli buffer at 95 °C for 5 min. (0.25–0.5 mm; ROTH). Equal protein concentration were loaded on 12% or 15% SDS–PAGE and transferred onto 0.45 µm nitrocellulose membranes (Amersham) using Tris-Glycine buffer supplemented with 10% ethanol. Membranes were blocked with TBST (20 mM Tris, 150 mM NaCl, pH 7.6 and 0.1% Tween20) with 5% non-fat dry milk for 30 min and incubated with anti-GFP primary antibody (1:5,000, Eurogentec 16B12), anti-Arf1 (1:5,000, Spang lab) or anti-Pgk1 primary antibody (1:5,000, Invitrogen clone 22C5D8) overnight at 4°C, followed by 2 h incubation with horseradish peroxidase (HRP)-conjugated secondary antibodies (1:10,000 anti-mouse, Invitrogen 31430 or 1:10,000 anti-rabbit, Sigma-Aldrich A0545) in TBST. Chemiluminescence signals were detected using Immobilon Western HRP Substrate (Millipore) and imaged using a FusionFX (Vilber Lourmat).

### Samples preparation for RNA and ribosomes purification

#### Subcellular Fractionation

Yeast cells from 2 L cultures were harvested and processed into frozen powder. The powder was resuspended in 500 μL cold extraction buffer (EP buffer: 20 mM HEPES/KOH pH 7.6, 100 mM sorbitol, 100 mM potassium acetate, 5 mM magnesium acetate, 1 mM EDTA, 100 μg/mL cycloheximide) supplemented with 1 mM DTT, 0.1 mM PMSF, and Halt protease inhibitor cocktail (Thermo Scientific). Following complete resuspension by vortexing, cellular debris was removed by centrifugation at 1,000 xg for 5 min at 4°C. The resulting supernatant was further centrifuged at 13,000 xg for 10 min at 4°C to yield a P13 pellet (membrane fraction) and an S13 supernatant (cytosolic fraction). The S13 fraction was subjected to an additional clarification spin at 20,000 xg for 15 min at 4°C to remove residual membrane components. The P13 pellet was washed once with EP buffer containing 1 mM DTT, 0.1 mM PMSF, protease inhibitors, and 2% polyoxyethylenlaurylether.

#### Sucrose Gradient Preparation

Linear 7-47% sucrose gradients were prepared using a gradient master (Biocomp). Sucrose solutions were prepared in APGB buffer (50 mM Tris/HCl pH 7.5, 50 mM NH₄Cl, 12 mM MgCl₂) supplemented with 0.5 mM DTT and 100 μg/mL cycloheximide. Gradients were prepared in polyethylene ultracentrifuge tubes. Gradients were allowed to equilibrate at 4°C for at least one hour before use.

#### RNase I Digestion and Ribosome Isolation

For ribosome footprinting, 10 A260 units of each fraction (P13 membrane or S13 cytosolic) were adjusted to 200 μL with EP buffer. Each sample was treated with 1.5 μL of RNase I (Ambion) and incubated for 1 h at 22°C with shaking at 1,400 rpm. The digestion was stopped by adding 10 μL of SUPERase•In RNase inhibitor (Ambion/Invitrogen). Samples were immediately loaded onto pre-cooled 7–47% linear sucrose gradients and centrifuged at 35,000 rpm for 2 h at 4°C in a TH-641/TST 41.14 rotor.

#### Gradient Fractionation

Gradients were fractionated using Gradient Master instrument (Biocomp) equipped with a UV detector set at sensitivity level 2. The pump speed was set to 0.75 mL/min, and fractions were collected every 32 sec (approximately 400 μL per fraction). Fractions were immediately flash-frozen in liquid nitrogen for subsequent RNA extraction and library preparation.

#### Total RNA Extraction

Total RNA was isolated from S13 and P13 fractions using phenol-chloroform extraction. Equal volumes of phenol-chloroform-isoamyl alcohol (PCI) mixture were added to each sample and vortexed for 10 seconds. Samples were centrifuged at 20,000 xg for 2 min at RT. The upper aqueous phase was transferred to fresh tubes, and the PCI extraction was repeated. RNA was precipitated by adding 0.1 volumes of 3 M NaCl and 2-3 volumes of 100% ethanol, followed by overnight precipitation at −80°C. Precipitated RNA was collected by centrifugation at 14,000 rpm for 30 min at 4°C. The RNA pellets were air-dried and resuspended in 30 μL of RNase-free water. RNA concentrations were determined using NanoDrop spectrophotometry.

#### mRNA Purification

Approximately 10 μg of total RNA was used for mRNA isolation using Dynabeads™ mRNA Purification Kit (Thermo Fisher Scientific) according to the manufacturer’s instructions. Purified mRNA was eluted in 20 μL of nuclease-free water, and 75 ng was used for subsequent custom RNA-Seq library preparation mimicking that of ribosome footprint.

#### RNA Fragmentation

mRNA were fragmented by alkaline hydrolysis by adding 50 μL of 2× alkaline hydrolysis buffer and incubated at 95°C for 5 min. Samples were then purified using RNeasy clean-up kit (Qiagen) and RNA was eluted in 18 μL of nuclease-free water.

#### RNA End Repair

Fragmented RNA were dephosphorylated in a 20 μL reaction containing 2 μL FastAP buffer, 1 μL RNasin, 1 μL FastAP enzyme, and 16 μL fragmented RNA. The reaction was incubated at 37°C for 30 min and heat-inactivated at 75°C for 10 min. Phosphorylation was performed by adding 5 μL of 10 mM ATP, 5 μL of PNK buffer A, 17 μL nuclease-free water, 1 μL RNasin, and 2 μL T4 PNK to the dephosphorylated RNA. The reaction was incubated at 37°C for 1 hour and purified using RNeasy columns, with elution in 16 μL nuclease-free water.

#### Adapter Ligation

For 3’ adapter ligation, 0.5 μL of the synthetized adapter RNA (universal Illumina adapters) was added to 14 μL of end-repaired RNA and incubated at 70°C for 2 minutes followed by immediate cooling on ice. The ligation reaction was completed by adding 1 μL RNasin, 2.5 μL T4 RNA ligase buffer (without ATP), 6 μL 50% DMSO, and 1 μL T4 RNA ligase (truncated). The reaction was incubated at 4°C overnight (18-20 h) and purified using RNeasy columns. For 5’ adapter ligation, 0.5 μL of the synthetized adapter RNA was added to 14 μL of 3’-ligated RNA and incubated at 70°C for 2 minutes followed by immediate cooling on ice. The ligation mixture was supplemented with 3 μL ligation buffer containing ATP, 9 μL 50% DMSO, 1 μL RNasin, 1 μL T4 RNA ligase, and 2 μL nuclease-free water. The reaction was incubated at 4°C overnight (minimum 12 h) and purified using RNeasy columns.

#### cDNA Synthesis and Amplification

Adapter-ligated RNA (10 μL) was mixed with 1 μL reverse transcription primer (100 μM), denatured at 72°C for 2 min, and immediately cooled on ice for 2 min. Reverse transcription was performed by adding 4 μL 5x First Strand buffer, 1 μL 10 mM dNTPs, 1 μL RNasin, 2 μL 0.1 M DTT, and 1 μL SuperScript III reverse transcriptase. The reaction was incubated at 48°C for 3 min, followed by 44°C for 60 min, and terminated at 75°C for 10 min. PCR amplification was performed using NEBNext master mix with 0.5 μL B222 forward primer (100 μM), 0.5 μL RPI index reverse primer (100 μM), and 2 μL cDNA in a final volume of 60 μL. The reaction was divided into three 20 μL aliquots and amplified using the following cycling conditions: initial denaturation at 98°C for 30 sec, followed by 10-14 cycles of 98°C for 10 sec, 65°C for 30 sec, and 72°C for 30 sec.

#### Sequencing and analysis

Samples were further sequenced with Single-reads 76 bases using the NextSeq 500 High Output Kit 75-cycles (Illumina, Cat# FC-404-1005)

#### Processing and analysis of RNA-seq data

Short single-end reads (63 nt) from RNA-sequencing (RNA-seq) samples (6 wile-type (WT) and 6 mutant (delARF1) samples) from supernatant fraction were equally processed with a customly developed snakemake^48^ pipeline using snakemake v.8.25.3. First, Illumina universal adapter sequences (starting with AGATCGGAAGAG) were trimmed from 3’ends of reads with cutadapt v.5.1^49^. Only reads with the length above 25 nt were retained. Reads were further mapped to the *S. cerevisiae* genome v.R64-1-1 with STAR aligner^50^ v.2.7.11b using the v.109 version of genome annotation. Genome and annotation files were downloaded from the Ensembl^51^ ftp site. STAR mapping was done with the following parameters: --runMode alignReads --alignEndsType Local --outSAMtype BAM SortedByCoordinate --outFilterType BySJout --outFilterMismatchNoverLmax 0.1 --outFilterMatchNmin 20 --alignIntronMin 10 -- alignSJoverhangMin 7 --alignSJDBoverhangMin 7 --outFilterScoreMinOverLread 0 --outFilterMatchNminOverLread 0 --twopassMode None --outBAMsortingThreadN {threads} --alignTranscriptsPerReadNmax 50000 --winAnchorMultimapNmax 50000 --outFilterMultimapNmax 50000 --outSAMprimaryFlag AllBestScore --outSAMattributes NH HI AS nM MD --limitOutSJcollapsed 5000000 --limitBAMsortRAM 15000000000 --limitOutSAMoneReadBytes 30000000. Uniquely mapped reads were used to quantify gene expression in strand-specific mode with featureCounts utility of subread package v.2.1.1^52^ using the following command-line parameters: featureCounts -O --fraction -s 1 -t exon -g gene_id.

#### Processing and analysis of Ribosome profiling (Ribo-seq) data

Short single-end reads (63 nt) from Ribo-seq samples (11 wile-type (WT) and 14 mutant (delARF1) samples) from the supernatant fraction were equally processed with a custom developed snakemake (v.8.25.3) pipeline^48^. Illumina universal adapter sequences (starting with AGATCGGAAGAG) and Ribo-seq-specific adapter sequences (starting with CTGTAGGCACCATCAAT) were trimmed from 3’ends of reads with cutadapt v.5.1^49^. Read length is a well-known confounding factor in Ribo-seq data analysis^53^, therefore remaining reads were separated by length, such that each read length in each sample defined an individual subsample. All such subsamples were further processed independently, in parallel. Only read lengths within the range 22-38 were allowed. The same reference genome as for RNA-seq data (see above), was used to map reads with STAR aligner^50^ v.2.7.11b, with the following parameters: --runMode alignReads --alignEndsType EndToEnd --runThreadN {threads} --genomeLoad NoSharedMemory --outSAMtype BAM SortedByCoordinate --outFilterType BySJout --outFilterMismatchNoverLmax 0.1 –outFilterMatchNmin 20 --outFilterScoreMinOverLread 0 --outFilterMatchNminOverLread 0 --twopassMode None -- alignTranscriptsPerReadNmax 50000 --winAnchorMultimapNmax 50000 --outFilterMultimapNmax 50000--outSAMprimaryFlag AllBestScore --outSAMattributes NH HI AS nM MD --limitOutSJcollapsed 5000000 --limitBAMsortRAM 15000000000 --limitOutSAMoneReadBytes 30000000. Secondary alignments were discarded, such that only multi-mapping (i.e. mapped to multiple genomic loci) reads with the same top alignment score for a read were retained. Rigorous quality control metrics were further obtained for reads of each read length in all the samples, inspired by and extending existing approaches^54–56^. First, the abundance of reads of each length relative to the total number of reads in each sample was calculated. Reads with lengths 22, 23, 37, and 38 had extremely low abundance across all samples and were discarded from further analysis. Second, the percentage of uniquely mapped reads among the reads used as input for the alignment step, was calculated. Third, proportions of all reads (both uniquely-mapped and multi-mapped) supporting expression of genes of different gene types (protein-coding, ribosomal RNA (rRNA), transfer RNA (tRNA), small nucleolar RNA (snoRNA) etc), were calculated using the featureCounts utility of the subread package v.2.1.1^52^ using the following command-line parameters: featureCounts -M -O --fraction -s 1 -t exon -g transcript_id. At that, a custom .gtf annotation file was prepared based on processed transcript Isoform sequencing (TIF-seq) data^57^ available at the Saccharomyces Genome Database (SGD) portal ^58^ and used as input. This annotation included untranslated transcribed sequences (UTRs). The file was prepared using python scripts and bedtools suite v2.31.1^59^ commands included in the supplementary jupyter notebook. Thus, the proportion of reads supporting expression of protein-coding genes, was used as a third quality metric. The fourth metric was the proportion of uniquely-mapped reads supporting expression from coding DNA sequence (CDS) relative to uniquely-mapped reads supporting CDS or UTR expression. Next, for each read length and every sample, the characteristic offset (nt distance) from 5′ end of the read to the first nucleotide of the P-site position in the translated mRNA was calculated^60^. For this, unique read alignments overlapping annotated start codons were extracted with a combination of bash and bedtools commands, and the relative proportions of reads supporting each of the possible valid P-site offsets (7-17 nt) was calculated. A single prominent offset is ideally expected, but in practice, several different offsets can have similar abundances. Therefore, a fourth quality metric was the percentage of reads supporting the most abundant (i.e. the most likely) offset, among the reads supporting any valid offset. Then, this offset was used as input to the custom python script to get single-nucleotide P-site positions across all genomic read alignments within a corresponding read length of a sample. Lastly, a combination of bedtools, bash and UCSC utilities^61^ were used to calculate the number of reads supporting every possible frame (0, +1, +2) of every codon in annotated CDS of protein-coding genes. The sixth quality metric was the relative proportion of reads supporting frame 0. All six quality metrics thus represented percentages in a range 0-100, and for each of them, higher values indicate better quality. Metrics were used together to cluster subsamples (sample and read length), using euclidean distance and the Ward hierarchical clustering approach.

#### Proteomics

Cells were grown overnight in YPD to reach mid-log phase, diluted to 0.2 OD600nm and grown for 6 h in fresh YPD, from which half the culture was shifted to 37°C for 30 min. 11×10^7^ cells were then spun down, washed twice in 10 mL 1x PBS and once in 1 mL of 1x PBS. Then cells were lysed at 4°C in 300 µL breaking buffer (25 mM Tris-HCl pH7.5, 150 mM NaCl, 2 mM EDTA, 0.6% Triton X100, 1 mM DTT, 1x Halt protease inhibitor cocktail (ThermoScientific)) supplemented with 100 µL of glass beads (0.25–0.5 mm; ROTH) for 20 min on a table-top multishaker. Cell debris and unbroken cells were pelleted by centrifugation 3,000 xg for 5 min at 4°C, incubated for 10 min at 95°C, alkylated in 20 mM iodoacetamide for 30 min at 25°C and proteins digested using S-Trap™ micro spin columns (Protifi) according to the manufacturer’s instructions. Briefly, 12 % phosphoric acid was added to each sample (final concentration of phosphoric acid 1.2%) followed by the addition of S-trap buffer (90% methanol, 100 mM TEAB pH 7.1) at a ratio of 6:1. Samples were mixed by vortexing and loaded onto S-trap columns by centrifugation at 4,000 xg for 1 min followed by three washes with S-trap buffer. Digestion buffer (50 mM TEAB pH 8.0) containing sequencing-grade modified trypsin (1/25, w/w; Promega, Madison, Wisconsin) was added to the S-trap column and incubate for 1 h at 47 °C. Peptides were eluted by the consecutive addition and collection by centrifugation at 4,000 xg for 1 min of 40 µL digestion buffer, 40 µL of 0.2% formic acid and finally 35 µL 50% acetonitrile, 0.2% formic acid. Samples were dried under vacuum and stored at −20 °C until further use. Peptides were labelled with isobaric tandem mass tags (TMTpro 16-plex, Thermo Fisher Scientific). After resuspension of the peptides in labeling buffer (2 M urea, 0.2 M HEPES, pH 8.3) by sonication, TMT reagents were added to the individual peptide samples followed by a 1 h incubation at 25°C shaking at 500 rpm. To quench the labelling reaction, aqueous 1.5 M hydroxylamine solution was added, and samples were incubated for another 5 min at 25°C shaking at 500 rpm followed by pooling of all samples. The pH of the sample pool was increased to 11.9 by adding 1 M phosphate buffer (pH 12) and incubated for 20 min at 25°C shaking at 500 rpm to remove TMT labels linked to peptide hydroxyl groups. Subsequently, the reaction was stopped by adding 2 M hydrochloric acid until a pH < 2 was reached. Finally, peptide samples were further acidified using 5% TFA, desalted using Sep-Pak Vac 1cc (50 mg) C18 cartridges (Waters) according to the manufacturer’s instructions and dried under vacuum.

TMT-labeled peptides were fractionated by high-pH reversed phase separation using a XBridge Peptide BEH C18 column (3.5 µm, 130 Å, 1 mm x 150 mm, Waters) on an Agilent 1260 Infinity HPLC system. Peptides were loaded on column in buffer A (20 mM ammonium formate in water, pH 10) and eluted using a two-step linear gradient from 2% to 10% in 5 min and then to 50% buffer B (20 mM ammonium formate in 90% acetonitrile, pH 10) over 55 min at a flow rate of 42 µL/min. Elution of peptides was monitored with a UV detector (215 nm, 254 nm) and a total of 36 fractions were collected, pooled into 12 fractions using a post-concatenation strategy as previously described^62^ and dried under vacuum.

Dried peptides were resuspended in 0.1% aqueous formic acid and subjected to LC–MS/MS analysis using a Q Exactive HF Mass Spectrometer fitted with an EASY-nLC 1000 (both Thermo Fisher Scientific) and a custom-made column heater set to 60°C. Peptides were resolved using a RP-HPLC column (75 µm × 30 cm) packed in-house with C18 resin (ReproSil-Pur C18–AQ, 1.9 μm resin; Dr. Maisch GmbH) at a flow rate of 0.2 µL/min. The following gradient was used for peptide separation: from 5% B to 15% B over 10 min to 30% B over 60 min to 45 % B over 20 min to 95% B over 2 min followed by 18 min at 95% B. Buffer A was 0.1% formic acid in water and buffer B was 80% acetonitrile, 0.1% formic acid in water.

The mass spectrometer was operated in DDA mode with a total cycle time of approximately 1 sec. Each MS1 scan was followed by high-collision-dissociation (HCD) of the 10 most abundant precursor ions with dynamic exclusion set to 30 sec. For MS1, AGC target was set to 3e6 with a maximum injection time of 100 ms and a resolution of 120,000 FWHM (at 200 m/z). For MS2 scans AGC target was set to 1e5 with a maximum injection time of 100 ms and a resolution of 30,000 FWHM (at 200 m/z). Singly charged ions and ions with unassigned charge state were excluded from triggering MS2 events. The normalized collision energy was set to 30%, the mass isolation window was set to 1.1 m/z and one microscan was acquired for each spectrum.

The acquired raw files were analysed using the SpectroMine software (Biognosis AG, Schlieren, Switzerland). Spectra were searched against a *Saccharomyces cerevisiae* database consisting of 6049 protein sequences (downloaded from Uniprot on 20190307) and 392 commonly observed contaminants. Standard Pulsar search settings for TMT 16 pro (“TMTpro_Quantification”) were used and resulting identifications and corresponding quantitative values were exported on the PSM level using the “Export Report” function. Acquired reporter ion intensities in the experiments were employed for automated quantification and statistical analysis using our in-house developed SafeQuant R script (v2.3)^63^. This analysis included adjustment of reporter ion intensities, global data normalization by equalizing the total reporter ion intensity across all channels, data imputation using the knn algorithm, summation of reporter ion intensities per protein and channel and calculation of protein abundance ratios. To meet additional assumptions (normality and homoscedasticity) underlying the use of linear regression models and t-tests, MS-intensity signals were transformed from the linear to the log-scale. The summarized protein expression values were used for statistical testing of between condition differentially abundant proteins. Here, empirical Bayes moderated t-tests were applied, as implemented in the R/Bioconductor limma package (http://bioconductor.org/packages/release/bioc/html/limma.html). The resulting per protein and condition comparison p-values were adjusted for multiple testing using the Benjamini-Hochberg method.

#### Affinity purification proteomics analysis

Following Arf1 immunoprecipitation, anti-GFP magnetic beads (Chromotek) were resuspended in elution buffer (5% SDS, 10 mM TCEP, 0.1 M TEAB), incubated for 10 min at 95°C and eluate was transferred into a new tube. Proteins were alkylated in 20 mM iodoacetamide for 30 min at 25°C and digested using S-Trap™ micro spin columns (Protifi) according to the manufacturer’s instructions. Briefly, 12% phosphoric acid was added to each sample (final concentration of phosphoric acid 1.2%) followed by the addition of S-trap buffer (90% methanol, 100 mM TEAB pH 7.1) at a ratio of 6:1. Samples were mixed by vortexing and loaded onto S-trap columns by centrifugation at 4,000 xg for 1 min followed by three washes with S-trap buffer. Digestion buffer (50 mM TEAB pH 8.0) containing sequencing-grade modified trypsin (1/25, w/w; Promega, Madison, Wisconsin) was added to the S-trap column and incubate for 1 h at 47 °C. Peptides were eluted by the consecutive addition and collection by centrifugation at 4,000 xg for 1 min of 40 µL digestion buffer, 40 µL of 0.2% formic acid and finally 35 µL 50% acetonitrile, 0.2% formic acid. Samples were dried under vacuum and stored at −20°C until further use.

Dried peptides were resuspended in 0.1% aqueous formic acid and subjected to LC–MS/MS analysis using a Orbitrap Fusion Lumos Mass Spectrometer fitted with an EASY-nLC 1200 (both Thermo Fisher Scientific) and a custom-made column heater set to 60°C. Peptides were resolved using a RP-HPLC column (75 µm × 36 cm) packed in-house with C18 resin (ReproSil-Pur C18–AQ, 1.9 µm resin; Dr. Maisch GmbH) at a flow rate of 0.2 µL/min. The following gradient was used for peptide separation: from 5% B to 12% B over 5 min to 35% B over 40 min to 50% B over 15 min to 95% B over 2 min followed by 18 min at 95% B. Buffer A was 0.1% formic acid in water and buffer B was 80% acetonitrile, 0.1% formic acid in water.

The mass spectrometer was operated in DDA mode with a cycle time of 3 sec between master scans. Each master scan was acquired in the Orbitrap at a resolution of 120,000 FWHM (at 200 m/z) and a scan range from 375 to 1600 m/z followed by MS2 scans of the most intense precursors in the linear ion trap at “Rapid” scan rate with isolation width of the quadrupole set to 1.4 m/z. Maximum ion injection time was set to 50 ms (MS1) and 35 ms (MS2) with an AGC target set to 1e6 and 1e4, respectively. Only peptides with charge state 2 – 5 were included in the analysis. Monoisotopic precursor selection (MIPS) was set to Peptide, and the Intensity Threshold was set to 5e3. Peptides were fragmented by HCD (Higher-energy collisional dissociation) with collision energy set to 35%, and one microscan was acquired for each spectrum. The dynamic exclusion duration was set to 30 sec.

The acquired raw-files were searched using MSFragger (v. 3.5) implemented in FragPipe (v. 18.0) against a *Saccharomyces cerevisiae* database (consisting of 6050 protein sequences downloaded from Uniprot on 20220222) and 392 commonly observed contaminants using the “LFQ-MBR” workflow. Quantitative data was exported from FragPipe and analyzed using the MSstats R package v.4.13.0^64^. Data was normalised using the default normalisation option “equalizedMedians”, imputed using “AFT model-based imputation” and p-values and q-values for pairwise comparisons were calculated as implemented in MSstats.

#### GO terms enrichment

Gene ontology terms of downregulated genes (RNAseq, RiboSeq) and decreased proteins (proteomics) were highlighted using the ShinyGo 0.77 website. Using the *Saccharomyces cerevisiae* database, an FDR cut-off of 0.05 was used and 20 pathways related to ‘Biological processes’ were shown with a pathway size of minimum 2 and maximum 2000 occurrences. Genes/proteins with a q-value >0.05 and a log2fold change of 1.5 were selected.

#### Statistics and reproducibility

All experiments were performed at least in three independent biological replicates. Unpaired two-tailed *t*-test or two-way analysis of variance (ANOVA) were calculated for each experiment using GraphPad Prism9. For image analysis, pictures were taken at random places on coverslips, analyzed in a blinded manner whenever possible with Omero web-client. All images are representative images from at least three independent biological experiments. For western blot, loading controls were run on a separate gel when the protein of interest had a similar molecular weight.

## Acknowledgments

We would like to thank Prof. Claudio de Virgilio for sharing the GFP-Tor1 integration plasmid; and Prof. Robbie Loewith for the generous gift of anti-Rps6 antibodies. We acknowledge the help and expertise of Sara Roig Romero from the Imaging core facility and Alex Schmidt from the Proteomics core facility of the Biozentrum and are grateful to Katharina Lobstein and Thomas Gross for technical assistance. Finally, we want to thank all members of the Spang lab for fruitful discussions.

## Funding

Swiss National Science Foundation: 310030B_163480, 310030_185127, 310030_219513, 320030-231859 University of Basel IdEx Unistra: ANR-10-IDEX-0002

## Author contributions

Conceptualization: LE, MZ, AS Investigation: LE, TN, AM, DR, CW Supervision: AS, MZ Writing—original draft: LE, AS Writing—review & editing: LE, TN, AM, DR, MZ, AS

## Competing interests

Authors declare that they have no competing interests.

## Data and materials availability

All data are available in the main text or the supplementary materials. RNAseq and Riboseq data are available at the NCBI SRA archive under the BioProject ID PRJNA1419792. Proteomics data are available at Massive UCSD website under MassIVE MSV000101167.

## Supplementary Materials for

**Fig. S1.**
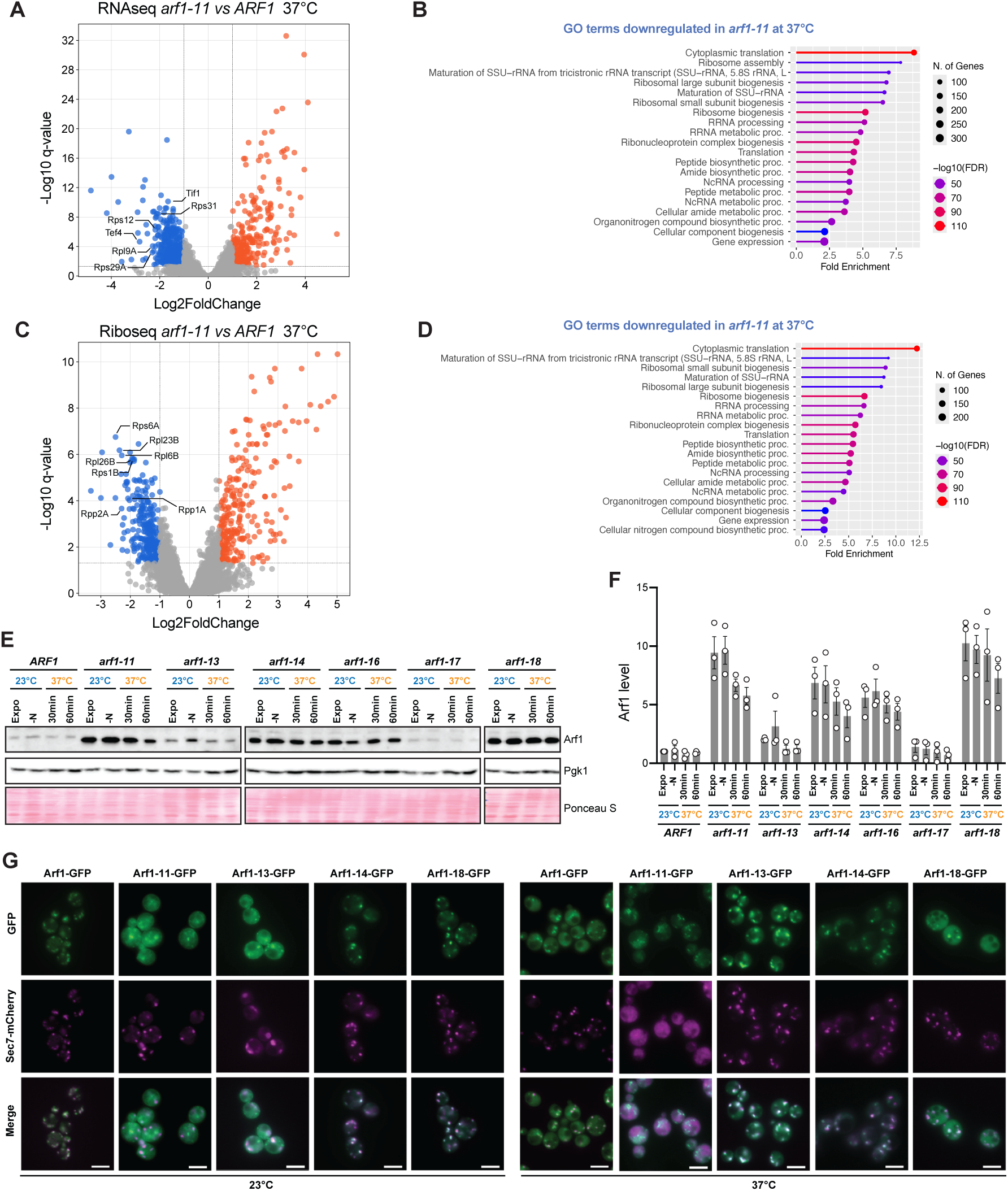
TORC1 activity is modulated by Arf1. **A)** Volcano plot obtained from *arf1-11 vs ARF1* strains by total RNAseq. All decreased transcripts in *arf1-11* at 37°C are shown in blue and increased transcripts in orange. A subset of decreased transcript related to ribosomes or translation regulation are shown. **B)** Gene Ontology terms associated to yeast biological processes based on decreased transcripts shown in (**A**). See also Supplementary Table 2. **C)** Volcano plot obtained from *arf1-11 vs ARF1* strains by Riboseq. All decreased transcripts in *arf1-11* at 37°C are shown in blue and increased transcripts in orange. A subset of decreased transcript related to ribosomes or translation regulation are shown. **D)** Gene Ontology terms associated to yeast biological processes based on decreased transcripts shown in (**C**). See also Supplementary Table 3. **E)** Arf1 immunoblot performed in 6 *ts*-allele mutants of *ARF1*, as well as WT *ARF1* strain. Pgk1 and membrane Ponceau S staining were used as loading controls. All strains were grown to exponential phase at 23°C for 6 h (Expo), shifted for 1 h into media lacking nitrogen sources (-N), or shifted for 30 or 60 minutes to 37°C. **F)** Arf1 protein levels measured based on (**E**). Mean and standard deviation are shown from n = 3 biological replicates. **G)** Widefield microscopy of Arf1 *ts*-alleles fused to GFP, expressing also Sec7-mCherry. Strains were grown to exponential phase at 23°C for 6 h or shifted for 30 minutes to 37°C. Scale bar: 5 µm.

**Fig. S2.**
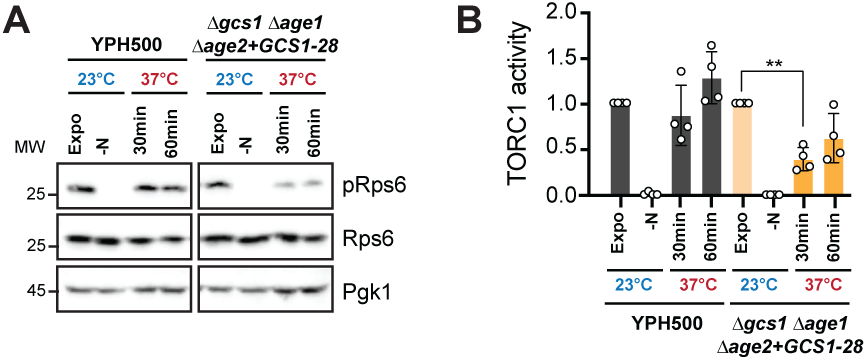
Arf1 activity regulates TORC1 activity. **A)** Immunoblot of Rps6 and its phosphorylated form pRps6 were performed in the YPH500 background strain, strains deprived of the ArfGEF *GCS1*, ArfGAP *AGE1* and *AGE2* rescued with the *ts*-allele *GCS1-28* (Δ*gcs1* Δ*age1 Δage2+ GCS1-28*). Pgk1 was used as loading control. Cells were grown to exponential phase at 23°C for 6 h (Expo), shifted for 1 h into media lacking nitrogen sources (-N), or shifted for 30 or 60 minutes to 37°C. **B)** Relative TORC1 activity based on protein level changes from immunoblots performed in (**D**). Mean and standard deviation are shown from n = 4 biological replicates. Two-way ANOVA with Turkey test Δ*gcs1* Δ*age1 Δage2 + GCS1-28* Expo *vs* Δ*gcs1* Δ*age1 Δage2 + GCS1-28* 30min \*\**P*: 0.0022.

**Fig. S3.**
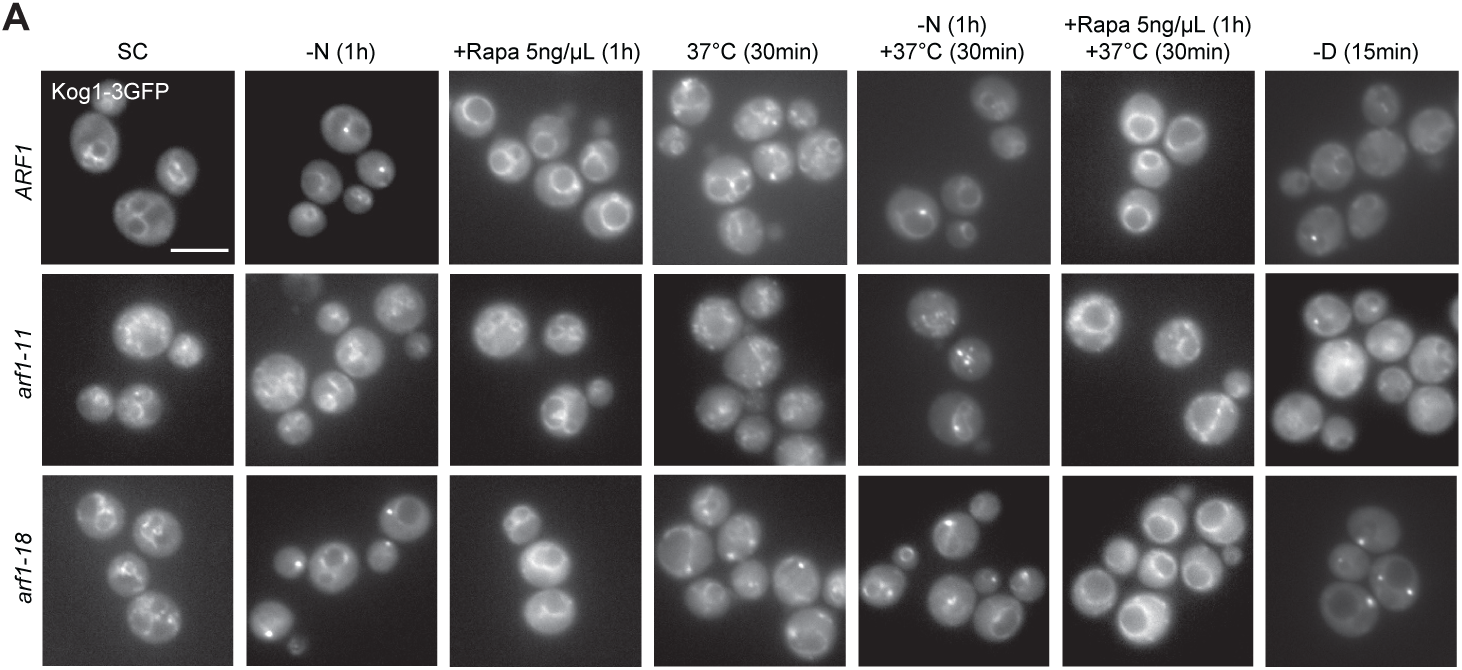
Arf1 interacts with TORC1 components in an allele-specific manner. **A)** Widefield microscopy of Kog1-3GFP expressed in *ARF1, arf1-11* and *arf1-18* strains. Strains were grown to exponential phase in synthetic media (SC), and shifted for 1h into media lacking nitrogen sources (-N), into SC media supplemented with Rapamycin (5 ng/mL), or shifted to 37°C for 30 minutes, shifted into -N media for 30 minutes at permissive temperature and then shifted for another 30 minutes to 37°C, into SC media supplemented with Rapamycin (5 ng/mL) for 30 minutes at 23°C and then shifted for another 30 minutes to 37°C, or shifted into SC media lacking glucose (-D) for 15 minutes. Scale bar: 5 µm.

**Fig. S4.**
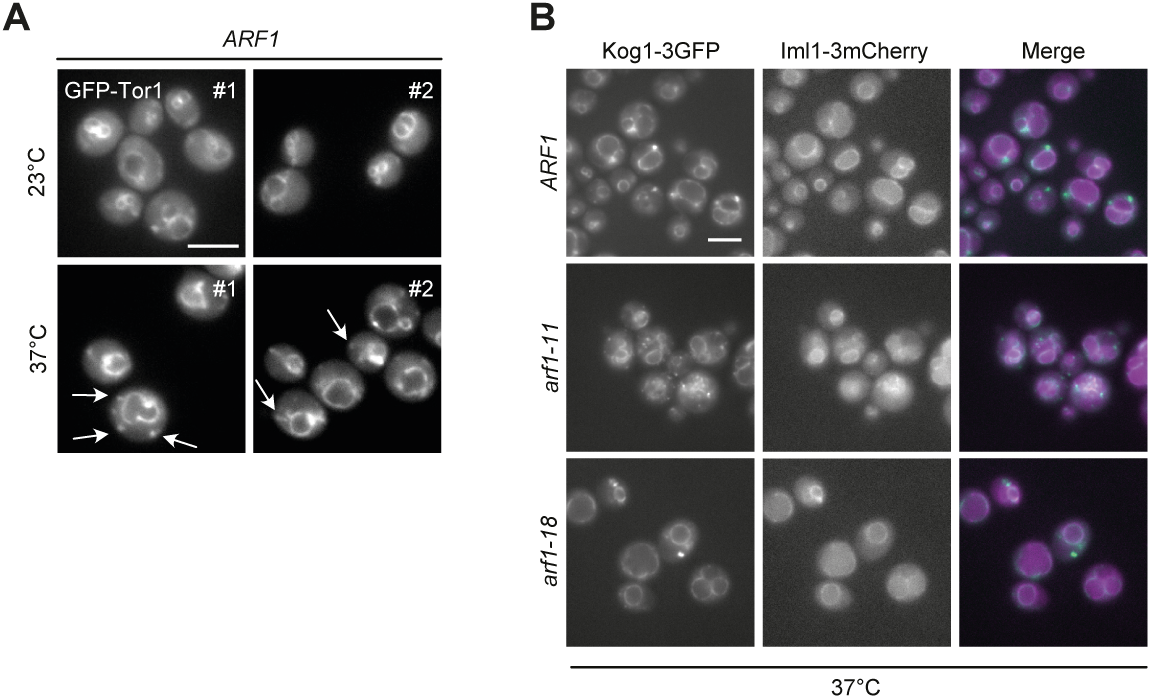
*arf1* mutants induce foci containing Kog1. **A)** Widefield microscopy of *ARF1* strain expressing Tor1 fused to GFP. The strain was grown to exponential phase at 23°C for 6 h or shifted for 30 minutes to 37°C. Arrows indicate GFP-Tor1 foci away from the vacuole. Two examples are shown for each condition. Scale bar: 5 µm. **B)** Widefield microscopy of *ARF1, arf1-11* and *arf1-18* strains expressing Kog1-3GFP and Iml1-3mCherry. The strain was grown to exponential phase at 23°C for 6 h or shifted for 30 minutes to 37°C. Scale bar: 5 µm.

**Fig. S5.**
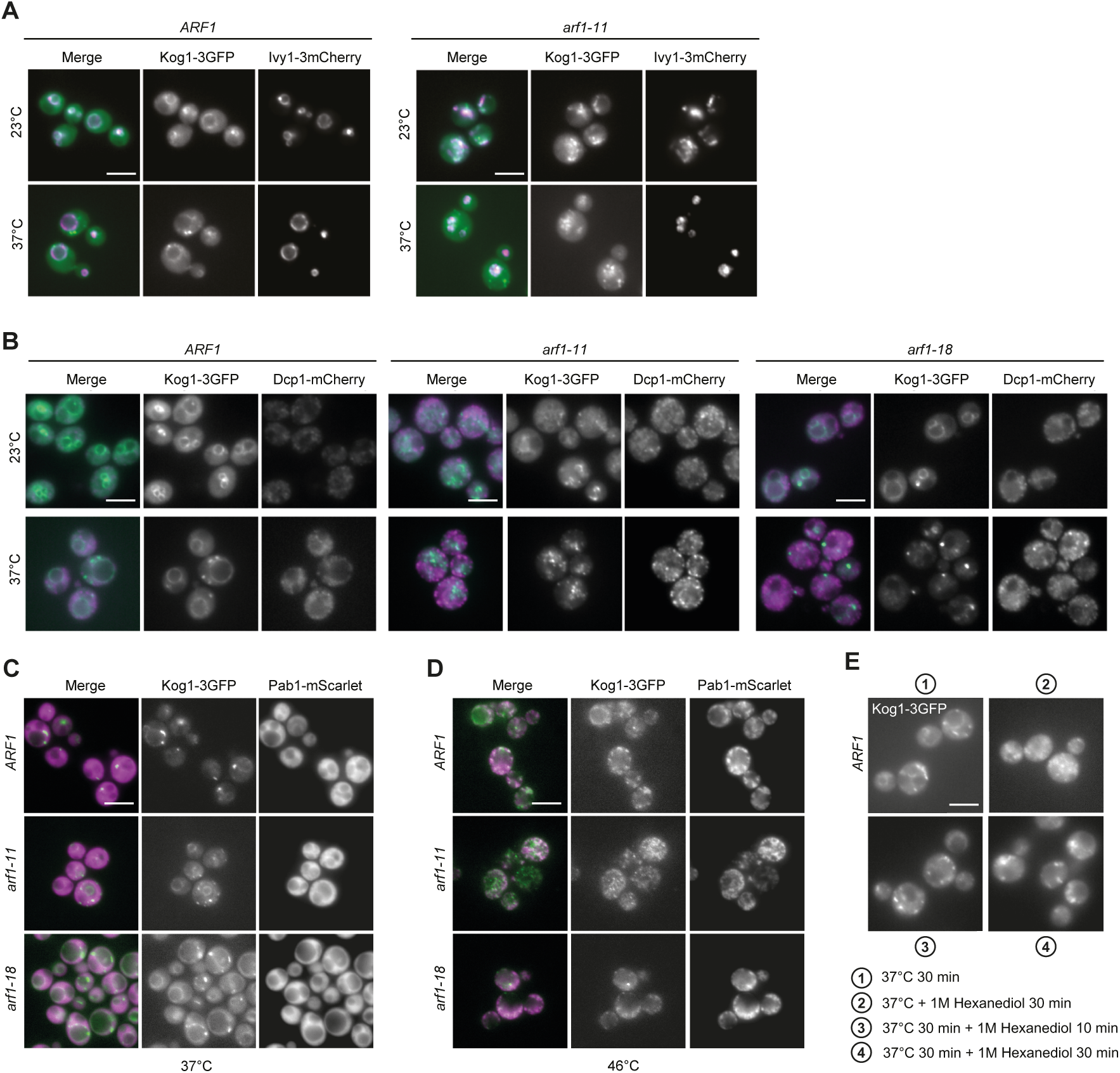
Kog1 foci represent a distinct reservoir for TORC1. **A)** Widefield microscopy of *ARF1* and *arf1-11* strains expressing Kog1-3GFP and Ivy1-3mCherry grown to exponential phase at 23°C and shifted for 30 minutes to 37°C. **B**) Widefield microscopy of *ARF1, arf1-11* and *arf1-18* strains expressing Kog1-3GFP and Dcp1-mCherry grown to exponential phase at 23°C and shifted for 30 minutes to 37°C. **C-D**) Widefield microscopy of *ARF1, arf1-11* and *arf1-18* strains expressing Kog1-3GFP and Pab1-mScarlet grown to exponential phase at 23°C and shifted for 30 minutes to 37°C (**C**) or 46°C (**D**). **E**) Widefield microscopy of *ARF1* strain expressing Kog1-3GFP grown to exponential phase at 23°C and shifted to 37°C for 30 minutes (1), shifted to 37°C and treated with 1 M 1,6-hexanediol for 30 minutes (2), or first shifted to 37°C for 30 minutes and then additionally treated with 1 M 1,6-hexanediol for 10 minutes (3) or 30 minutes (4). Scale bar in all panels: 5 µm.

**Fig. S6.**
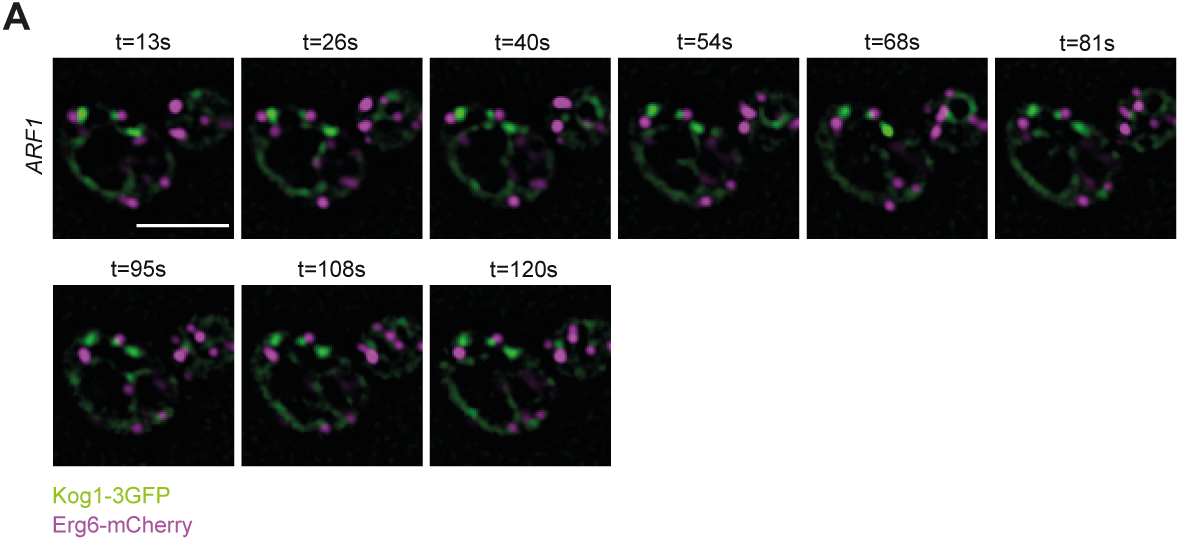
ER-anchored Arf1-11 partially rescues TORC1 activity at the restrictive temperature. **A)** Widefield microscopy of *ARF1* strain expressing Kog1-3GFP and Erg6-Cherry grown to exponential phase at 23°C and shifted for 30 minutes to 37°C. Time stamps are representative images based on 120 seconds movies. Images were processed by Gaussian blur in Fiji. Scale bar: 5 µm.

---

**Table S1.** *arf1-11* vs *ARF1* proteome data from mass spectrometry analysis (related to Fig. 1)

**Table S2.** Transcripts from total RNA sequencing analysis (related to Fig. S1)

**Table S3.** Transcripts from ribosome footprint sequencing analysis (related to Fig. S1)

**Table S4.** Arf1-11-GFP vs Arf1-GFP immunoprecipitation data from mass spectrometry analysis (related to **Fig. 3**)

## Notes

### Competing Interest Statement

The authors have declared no competing interest.

